# Beyond deadwood volume: structural heterogeneity and species-specific dispersal shape saproxylic beetle genetics

**DOI:** 10.64898/2026.07.17.738652

**Authors:** Rama Sarvani Krovi, Nermeen R. Amer, Alicja Wierzbicka, Radosław Plewa, Marcin Kadej, Tomasz Jaworski, Adrian Smolis, Tomasz Szmatoła, Maria Oczkowicz, Łukasz Kajtoch

**Author notes:** Corresponding author: Rama Sarvani Krovi. joint first authors (Rama Sarvani Krovi, Nermeen R. Amer).

## Abstract

Many rare saproxylic (deadwood-dwelling) beetles are among Europe’s most threatened insects, yet the relative importance of local microhabitat quality versus broad-scale landscape connectivity in shaping their genetic integrity remains poorly understood. We evaluated environmental drivers for 13 saproxylic beetle species—rare old-growth specialists, common taxa, and economically significant pests—across eight major forest complexes in Poland, using double digest restriction-site associated DNA sequencing (ddRAD-seq) paired with plot-level structural inventories and estimated effective migration surfaces (EEMS) modelling. A beta generalized linear mixed model identified deadwood diversity as the only significant within-species predictor of observed heterozygosity (β = 0.041, 95% CI 0.0227–0.0586, p < 10⁻⁵); total deadwood volume, forest age, time since logging, stump density, and veteran tree density were not significant. Management-category contrasts in observed heterozygosity (Ho) and the inbreeding coefficient (Fis) were non-significant after Tukey adjustment. Predictor x ecological-tier interactions did not differ detectably between rare and common species for five of six structural predictors; time since last logging was the exception (χ² = 4.56, p = 0.033). EEMS patterns differed among species and fell into three broad groups: regional lineage barriers, asymmetrical corridors, and surfaces close to isolation-by-distance expectations. These results associate plot-level heterozygosity with deadwood diversity and show that regional gene-flow patterns cannot be generalised across saproxylic beetles. Conservation planning should therefore combine local deadwood restoration with species-specific assessment of landscape connectivity.

## Introduction

Saproxylic (i.e. deadwood-dwelling) beetles are ecologically critical components of forest systems, mediating nutrient cycling, organic decomposition, and trophic dynamics (Siitonen 2001). Europe harbours at least approximately 1,500 currently recognised saproxylic beetle species (Gutowski et al. 2023), representing approximately 5% of the regional Coleoptera fauna (Slipinski et al. 2011); this figure is likely a substantial underestimate when facultatively saproxylic taxa are included (Gimmel & Ferro 2018). Their ecological importance is matched by a severe conservation crisis: nearly 700 species appear on the IUCN European Red List, with many reliant on old-growth structural features that are increasingly absent from managed landscapes (Calix et al. 2018).

The persistence of saproxylic communities is tightly linked to the spatial and temporal continuity of coarse woody debris and cavity-bearing veteran trees (Lachat et al. 2013; Müller et al. 2015). Primeval forests may contain deadwood volumes of 250–300 m³/ha (Puletti et al. 2017; Oettel et al. 2023), whereas intensively managed commercial forests frequently contain less than 5 m³/ha (Bobiec & Stachura-Skierczyńska 2007; Godzik & Piechnik 2019). Müller and Bütler (2010) reported ecological thresholds ranging from 10 to 150 m³/ha, including estimates of 20–30 m³/ha for boreal coniferous forests, 30–40 m³/ha for mixed-montane forests, and 30–50 m³/ha for lowland oak–beech ecosystems. Many intensively managed forests therefore contain less deadwood than the ranges proposed for vulnerable deadwood-dependent communities.

Habitat fragmentation and resource scarcity can affect genetic variation before demographic declines are detected. Reported effects include reduced effective population size (Ne), loss of allelic richness (Ar), lower observed and expected heterozygosity (Ho and He), and increased local inbreeding (Fis) (Woodruff 2001; Davies et al. 2008). For species with limited dispersal, logging of mature trees and removal of deadwood can reduce habitat continuity and gene flow (Oleksa et al. 2015; Henneberg et al. 2021, 2025). Isolation-by-distance models have commonly been used to describe these patterns, whereas genomic data allow spatial modelling of migration corridors and barriers (Cole et al. 2020). In Central European saproxylic beetles, regional genetic structure has been associated with forest naturalness, and primeval forest remnants may retain genetic variation lost elsewhere (Kajtoch et al. 2022). These findings suggest that both local habitat structure and the regional distribution of suitable forest can influence gene flow.

Comparative assessments across management regimes and broad geographic regions remain scarce (Komonen & Müller 2018; Kajtoch et al. 2022). Landscape metrics alone may miss associations between genetic variation and local structural variables such as deadwood and forest age, whereas plot-level data do not describe regional migration. Combining multilocus genomic data with structural inventories allows both scales to be examined in the same analysis (Rellstab et al. 2015).

In this study, we evaluated the contributions of local forest structure and regional landscape connectivity to the genetic integrity of 13 saproxylic beetle species across Poland, Central Europe. These species represent distinct ecological strategies and conservation statuses, including rare old-growth specialists, common forest taxa, and economically significant forest pests. We tested the following hypotheses:

H1: Plot-level structural integrity (total deadwood volume, deadwood diversity, forest age, time since logging, stump density, and veteran tree density) predicts within-population genetic diversity (Ho) and inbreeding (Fis), with structurally rich forests associated with higher diversity and lower inbreeding.

H2: Effective migration surfaces differ among ecological strategies, with rare old-growth specialists expected to show stronger or more spatially restricted barriers than broadly distributed common and pest taxa.

H3: Rare and common saproxylic beetle species respond congruently to the same environmental gradients—that is, the direction and approximate magnitude of structural effects on genetic diversity are shared between ecological tiers.

## Methods

### Study Area and Sampling Design

We sampled saproxylic beetles across eight major forest complexes in Poland, partitioned into three macro-regions: (1) Northeast Lowlands: Białowieża Forest, Knyszyn Forest, and Augustów Forest; (2) South-Central and Southeast Mountains: the Holy Cross Mountains and the Carpathians; and (3) Southwest Lowlands: Lower Silesia Forest, Barycz Valley Forest, and Oder Valley Forest. Sampling sites represented four forest management and protection categories: primeval-protected (national parks or nature reserves), reserves (nature reserves within commercial forests), primeval-non-protected (primeval forests outside protected areas), and commercial (logged forests without formal protection). This design incorporated all three remaining primeval forest complexes in Poland: Białowieża Forest, the Carpathian forests, and the Holy Cross Mountains (Figure 1; Supplementary Table S1). The sampling design covered contrasting management and protection regimes and provided sufficient numbers of sites and individuals for statistical analysis.

**Figure 1.**
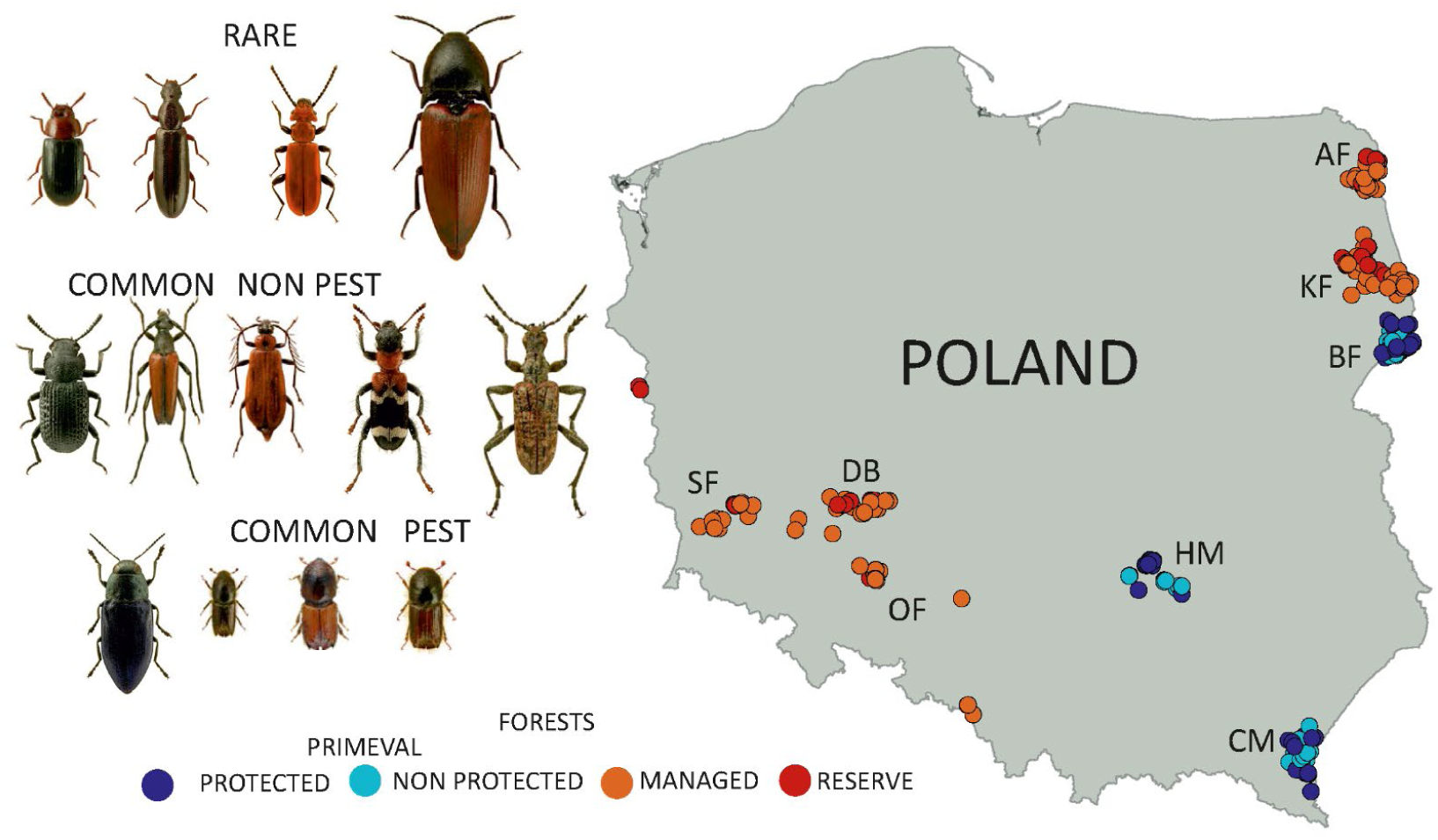
Species illustrations reproduced from Iconographia Coleopterorum Poloniae © Lech Borowiec, used with permission. Sampling locations across Poland considered in the study classified by ecological tier. Species (from left to right): Rare: *Neomida haemorrhoidalis, Boros schneideri*, *Cucujus cinnaberinus*, *Elater ferrugineus*; Common: *Bolitophagus reticulatus*, *Schizotus pectinicornis*, *Stenurella melanura*, *Thanasimus formicarius, Rhagium inquisitor*; Pest: *Ips acuminatus*, *Ips sexdentatus*, *Ips typographus*, *Phaenops cyanea*. Forest labels: AF - Augustów Forest, KF - Knyszyn Forest, BF - Białowieża Forest, HM - Holy Cross Mountains, CM - Carpathians, OF - Oder Forest, DB - Barycz Forest, SF - Silesia Forest.

Beetle collection was conducted during 2022–2024. Of 442 initially selected sites, 430 were retained for ecological and genomic analysis. We applied two collection methods: (1) hand collecting for majority of species involving direct microhabitat searches beneath bark, within decayed wood, inside tree cavities, and inside sporocarps of bracket fungi for *Bolitophagus reticulatus* and *Neomida haemorrhoidalis*; and (2) non-destructive collection using pheromone-baited barrier traps. The latter method was used for cryptic taxa, including *Ips* species, and for *Elater ferrugineus* (Bouget et al. 2008; Galko et al. 2016; Kärvemo et al. 2025). All collected specimens were immediately preserved in 96% ethanol and stored at −20 °C before DNA extraction (Reiss et al. 1995). From approximately 250 captured saproxylic species, genomic analysis was performed for 13 target species represented at a minimum of 20 independent geographic sites across multiple forest types. We categorised these species into three ecological tiers: rare/relict stenotopic species - *Boros schneideri* (Panzer, 1796), *Cucujus cinnaberinus* (Scopoli, 1763), *Neomida haemorrhoidalis* (Fabricius, 1787), and *Elater ferrugineus* (Linnaeus, 1758); common species - *Thanasimus formicarius* (Linnaeus, 1758), *Bolitophagus reticulatus* (Linnaeus, 1767), *Schizotus pectinicornis* (Linnaeus, 1758), *Stenurella melanura* (Linnaeus, 1758), and *Rhagium inquisitor* (Linnaeus, 1758); and pest species - *Ips acuminatus* (Gyllenhal, 1827), *Ips sexdentatus* (Börner, 1776), *Ips typographus* (Linnaeus, 1758), and *Phaenops cyanea* (Fabricius, 1775). To balance DNA yield with voucher preservation, handling protocols varied according to taxon size and conservation status.

### Multi-Scale Ecological Data Collection

Each study site was georeferenced using a high-precision GPS device. At each site, we established a 500 m² circular plot (radius 12.6 m) for structural inventory. Deadwood was quantified as the total volume of coarse deadwood, excluding twigs and roots, categorised by orientation (standing or lying) and wood type (coniferous or deciduous). A deadwood diversity index (Shannon-equivalent diversity across decay class x wood type categories) was calculated for each plot. Microhabitat metrics included dead tree species richness, old and fresh stump densities, and percentage dead branch cover on the forest floor. Stumps were classified as old or fresh from cut-surface weathering and wood decay; old stump density was used in plot-level models, whereas fresh stump density was retained for descriptive summaries. Veteran trees were defined as living monumental trees with advanced microhabitat features, including cavities, fungal fruiting bodies, and large dead branches, and were recorded using diameter-at-breast-height thresholds of >100 cm for oak and beech and >80 cm for all other tree species. Macrostructural variables were extracted from the Polish Forest Data Bank (BDL; https://www.bdl.lasy.gov.pl/portal): canopy cover, understorey cover, forest age (the age of the dominant tree species), total volume of living trees, and tree species richness. Time since the most recent intensive management activity, such as logging or thinning, was determined from protection-designation dates for national parks and strict reserves; for commercial forests, it was derived from BDL harvesting records and corroborated by field assessment of the ratio of highly decayed to freshly cut stumps. Surrounding forest cover was calculated within concentric buffers of 100 and 5000 m using CORINE Land Cover 2018 data in QGIS 3.34 (QGIS Association 2023). The environmental characteristics of all sampling sites are summarized in Supplementary Table S1 and S2. To avoid model overparameterisation and maintain a clear distinction between spatial scales, environmental variables were partitioned before analysis. Plot-level models used six direct management and deadwood indicators: total deadwood volume, deadwood diversity, forest age, time since logging, old stump density, and veteran tree density.

### Laboratory Protocol and ddRAD-seq Library Preparation

Genomic DNA was extracted using the Sherlock AX magnetic-bead kit (A&A Biotechnology, Poland). Tissue input was optimised by species size: whole-body tissue for the smallest species, head-and-thorax fragments for medium-sized species, and single legs for the largest, strictly protected taxa due to permission constraints. DdRAD- seq libraries were prepared following Bayona-Vásquez et al. (2019) using NheI and EcoRI restriction enzymes. Individual fragments were ligated with internally barcoded adapters and size-selected to 400-600 bp. Multiplexed pools were sequenced on an Illumina NovaSeq X Plus platform generating 150-bp paired-end reads targeting an average depth of 4 million raw reads per individual at Admera Health (NJ, USA).

### Bioinformatics and population genetic analyses

Raw reads were quality filtered, checked for adapter contamination, and demultiplexed in Stacks v2.68 (Rochette et al. 2019); reads with Phred scores <30 or uncalled bases were removed. Because reference genomes were unavailable for most target taxa, locus assembly was performed de novo. Assembly parameters were a minimum stack depth of 3 (-m 3) and a species-specific inner-locus mismatch tolerance of M = 2. The final single-nucleotide-polymorphism (SNP) dataset retained loci present in at least 80% of individuals within each population (-r 0.80), with a minimum minor-allele-frequency threshold of 0.03 and a minimum read depth of 5x per genotype. Only the first verified SNP per RAD locus was exported to avoid inflating linkage disequilibrium. Per-population observed heterozygosity (Ho) and the inbreeding coefficient (Fis) were calculated with the R packages hierfstat (Goudet 2005) and adegenet (Jombart 2008). Deviations from Hardy–Weinberg equilibrium were evaluated by testing Fis significance with 10,000 permutations. The population-level dataset, including coordinates, management category, Ho, He, Fis, and allelic richness, is provided in Supplementary Table S1.

### Statistical Modelling

#### Forest structural predictors of genetic diversity (H1)

To test whether plot-level forest structure predicts genetic diversity and inbreeding (H1), we used a multistage modelling approach. First, multivariate differences among management categories were tested by PERMANOVA on Euclidean distances calculated from six z-scored structural variables: total deadwood volume [V_total], deadwood diversity [deadwood_diversity], forest age [max_age], time since logging [Latest_logging], old stump density [stumps_old], and veteran tree density [veterans]. The analysis used vegan::adonis2 (Oksanen et al. 2025; Anderson & Walsh 2013) with 9,999 permutations under a reduced model; vegan::betadisper was used to assess multivariate dispersion. We then modelled observed heterozygosity (Ho, bounded 0–1) and the inbreeding coefficient (Fis). For Ho, we fitted a generalized linear mixed model (GLMM) with a beta error distribution and logit link via glmmTMB (Brooks et al. 2017). Fixed effects were the management type (primeval-protected, primeval-non-protected, reserves, and commercial) and the six standardised structural predictors. Site was included as a random intercept, and genus and species were included as nested random intercepts (1 | Genus / Species) to account for taxonomic relatedness. An analogous linear mixed model with a Gaussian error structure was fitted for Fis.

Tukey HSD post-hoc contrasts among management categories were obtained from the fitted models (Agbangba et al. 2024). Marginal (R²m) and conditional (R²c) coefficients of determination were computed following Nakagawa et al. (2017) with the MuMIn package (Bartoń 2010). Residual spatial autocorrelation was assessed post-hoc using Moran’s I on residuals aggregated by unique sampling locations via DHARMa::testSpatialAutocorrelation (Hartig 2022). To obtain species-resolved effect sizes, per-species linear regressions of Ho against each structural predictor (Ho ∼ scale(predictor)) were fitted with the metafor package (Viechtbauer 2010), with p values corrected across the species x predictor matrix using the Benjamini–Hochberg false-discovery rate (Benjamini & Hochberg 1995).

#### Effective migration surfaces among ecological strategies (H2)

To evaluate spatial genetic structure and effective migration rates, we reconstructed spatial gene-flow dynamics using estimated effective migration surfaces (EEMS; Petkova et al. 2016). EEMS maps spatial deviations from standard isolation-by-distance expectations by projecting georeferenced population-genetic data onto a dense grid of demes and estimating relative effective migration rates (m) across the landscape. Separate EEMS models were run for each of the 13 species with a spatial grid of 200 demes. Model fit was assessed by comparing observed and fitted between-deme dissimilarities (Supplementary Figure S1). Migration surfaces were optimised using Markov chain Monte Carlo sampling for 2,000,000 iterations, with 50,000 discarded as burn-in and every 10,000th iteration retained. Three independent chains per species, initiated with different random seeds, were used to assess convergence. Convergence was evaluated by visually inspecting trace plots and monitoring log-likelihood values across iterations. Posterior median migration rates and their uncertainty were visualised with rEEMSplots.

#### Environmental-response congruence across ecological tiers (H3)

To evaluate whether rare and common saproxylic beetle species respond congruently to environmental gradients (H3), secondary beta GLMMs for Ho and Gaussian mixed models for Fis were fitted to 312 site-level population records from nine rare and common species. Pest species were excluded because H3 specifically compares the two conservation-relevant ecological tiers. Models included the six structural predictors, ecological tier, and each predictor x ecological-tier interaction. Type-II Wald χ² tests of interaction terms provided the formal test of H3: a non-significant interaction indicated that response slopes did not differ detectably between tiers. Tier-specific slopes were extracted with emmeans::emtrends. Baseline tier differences in Ho and Fis were assessed with Wilcoxon rank-sum tests and Benjamini–Hochberg adjustment.

## Results

### Characteristics of study sites

This study assembled forest structural data from managed, reserve, and primeval forests across Poland by combining field measurements of deadwood and microhabitats with stand attributes from the Polish Forest Data Bank (Supplementary Table S1). Median dominant tree age was 140 years in primeval-protected sites and reserves, 97 years in primeval-non-protected forests, and 87 years in commercial forests. The median volume of living trees was approximately 440 m³/ha in reserves and 425 m³/ha in primeval-protected sites, compared with 325 and 323 m³/ha in primeval-non-protected and commercial forests, respectively. Tree species richness was similar across categories (median 6–7 species). Time since the last logging strongly differentiated forest categories: medians were 34–38 years in reserves and primeval-protected sites and ≤5 years in commercial and primeval- non-protected forests. Old stump density, a proxy for cumulative harvest intensity, had a median of 0 per plot in reserves and primeval-protected sites and 3.0 in commercial forests; fresh stumps had a median of 0 across categories. Median total deadwood volume was approximately 150 m³/ha in reserves and primeval-protected sites, 88 m³/ha in primeval-non-protected forests, and 31 m³/ha in commercial forests. Deadwood diversity showed a similar contrast, with a median of 2 in the protected categories and 1 in commercial forests (Table 1). Dead branch cover was broadly similar among categories (median 20–30%), whereas veteran tree density declined from a median of 1.0 per plot in primeval-protected sites to 0.0 in commercial forests. Canopy, understorey, and shrub-layer cover and forest cover within the defined buffer zones (Supplementary Table S1 and S2) did not differ significantly among forest types. Subsequent analyses therefore focused on the six a priori structural predictors most directly related to saproxylic resources and management history.

**Table 1.** Mean structural and compositional values for forest plots categorised by protection and management status. Per-plot deadwood values are in m³ per 500 m² plot (multiply by 20 for m³/ha-equivalent). Stump density, veteran tree density, and time since latest logging are summarised in the Results and given in full in Supplementary Table S1; they are omitted here to keep the table concise.

| Structural variables | Commercial<br>(n=160) | Reserves<br>(n=84) | Primeval-non-protected<br>(n=97) | Primeval-protected<br>(n=89) | Overall<br>(n=430) |
| --- | --- | --- | --- | --- | --- |
| Max_age (year) | 84.2 ± 41.8 | 130.3 ± 47.2 | 107.8 ± 40.9 | 136.0 ± 42.6 | 109.3 ± 47.9 |
| Total deadwood volume (m<sup>3</sup>/ha) | 70 ± 96 | 215 ± 241 | 120 ± 110 | 216 ± 171 | 140 ± 166 |
| Deadwood diversity (Shannon)_ | 1.41 ± 0.86 | 2.14 ± 0.78 | 1.71 ± 0.80 | 1.83 ± 0.74 | 1.71 ± 0.85 |
| Latest_Logging (year) | 3.4 ± 3.0 | 45.0 ± 15.9 | 7.6 ± 10.5 | 43.9 ± 24.0 | 20.8 ± 23.9 |
| Stumps_old (N per plot) | 4.79 ± 4.81 | 1.74 ± 3.69 | 1.87 ± 3.10 | 0.94 ± 1.46 | 2.74 ± 4.05 |
| Veterans (N per plot) | 0.38 ± 0.86 | 1.15 ± 1.68 | 0.90 ± 1.42 | 1.31 ± 1.34 | 0.84 ± 1.34 |

Forest age, volume of living trees, total deadwood volume, old stump density, veteran tree abundance, and deadwood composition varied among study sites, whereas tree species richness was comparatively similar (Table 1). Rare specialists (*B. schneideri*, *C. cinnaberinus*, *E. ferrugineus*, and *N. haemorrhoidalis*) were recorded mainly at sites with an average total deadwood volume of 160–180 m³/ha and were difficult to collect in commercial forests. *C. cinnaberinus*, *E. ferrugineus*, and *S. pectinicornis* were associated with decayed logs. *E. ferrugineus* and *N. haemorrhoidalis* were associated with deciduous deadwood and occurred at sites with the highest veteran tree counts, whereas *B. schneideri* was associated with standing coniferous deadwood averaging 134 m³/ha. Pest species showed the opposite pattern: *P. cyanea* and *I. acuminatus* occurred mainly in commercial forests containing less than 40 m³/ha of deadwood, predominantly coniferous slash. Common taxa and *I. typographus* occupied sites with intermediate deadwood volumes (60–100 m³/ha). Here, deadwood composition refers specifically to deadwood diversity and the representation of standing versus downed and coniferous versus deciduous substrates.

Forest management categories differed in their multivariate structural profiles (PERMANOVA: pseudo F_3426_ = 41.55, R² = 0.226, p < 0.001; Figure 2; Table 2). PCo1 and PCo2 accounted for 37.4% and 16.3% of the ordination variation, respectively. Dispersion also differed among categories (PERMDISP: F_3426_ = 5.83, p = 6.6 x 10⁻⁴). Because PERMANOVA is sensitive to unequal dispersion, the significant result may reflect both differences in average structural composition and differences in variability within categories. It is interpreted here as evidence of structural differentiation, not as proof that management category caused the separation.

**Figure 2.**
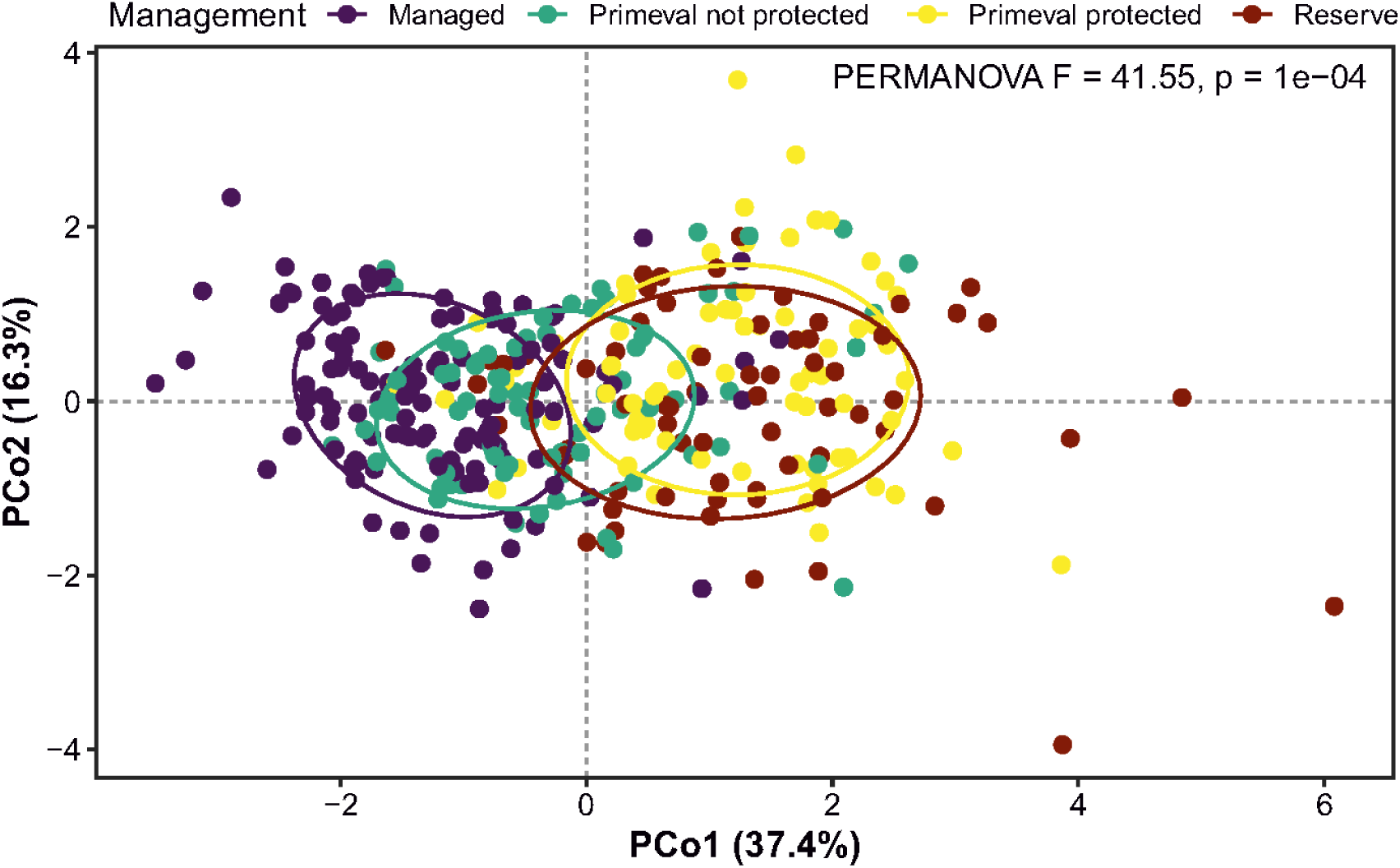
Principal coordinates ordination (PCoA) of forest structural variables across 430 sampling plots, coloured by management type.

**Figure 3.**
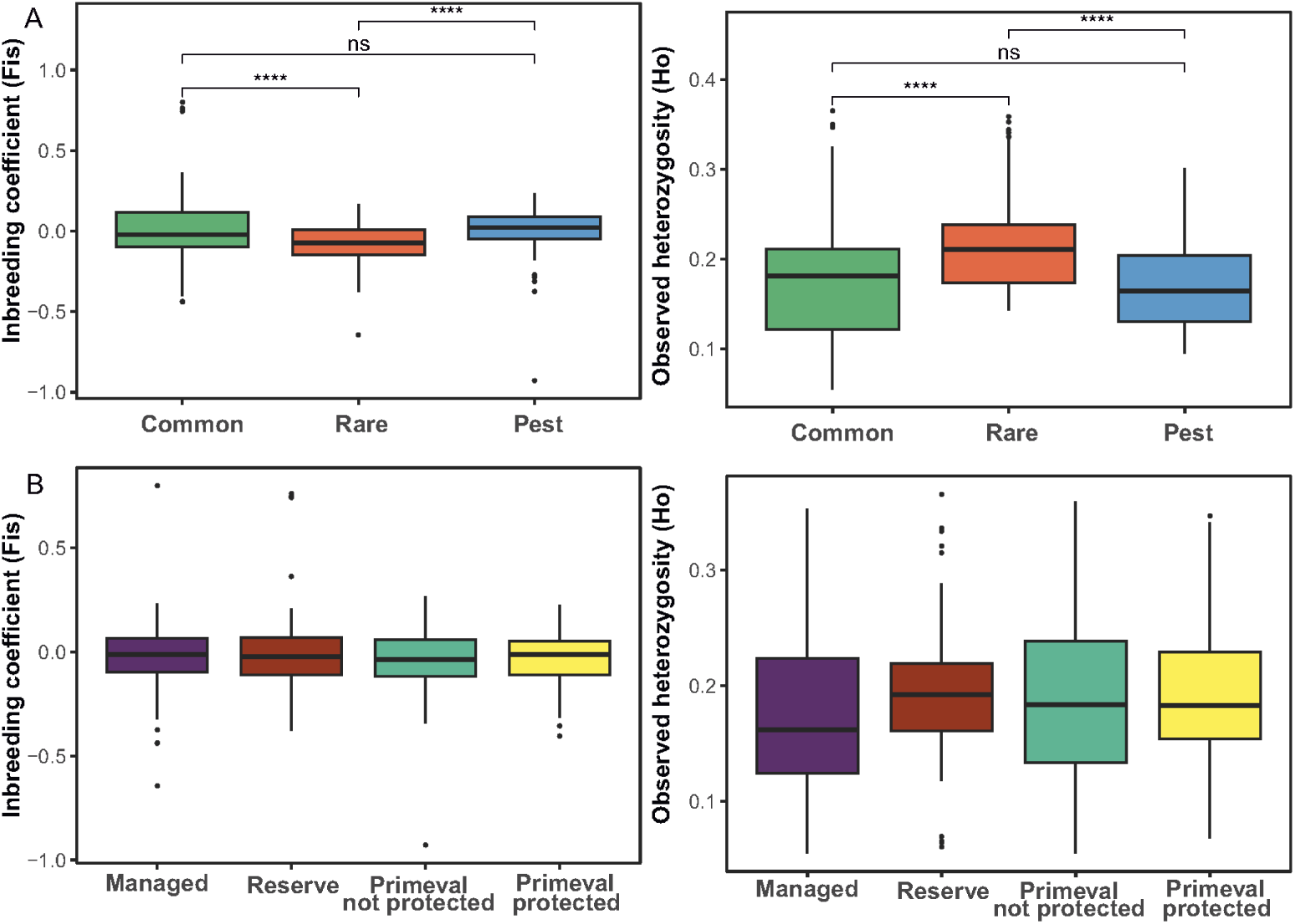
Boxplots displaying differences in observed heterozygosity (Ho) and inbreeding coefficient (Fis) across ecological tiers (Common = green, Rare = red, Pest = blue) and management categories (lower panel). Tier comparisons by Wilcoxon rank-sum tests with Benjamini-Hochberg correction; **** denotes p < 10⁻⁴.

**Table 2.**
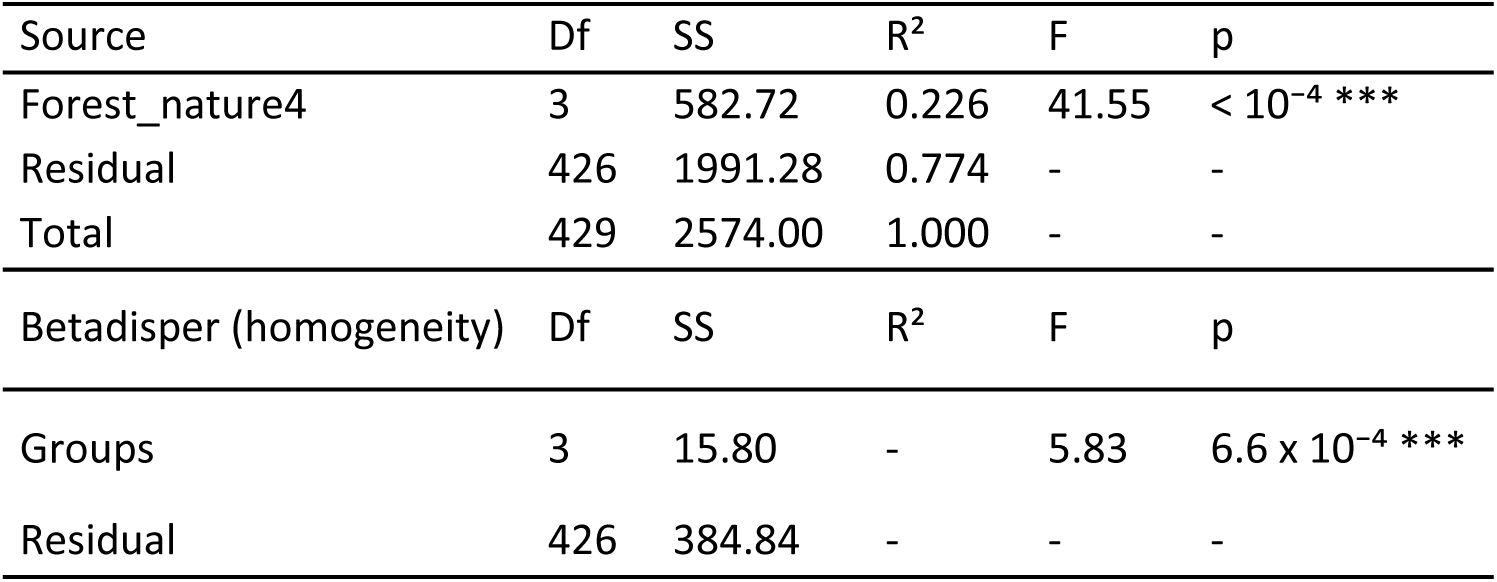
PERMANOVA and dispersion-homogeneity tests for forest structural variables across management categories (Euclidean distance, 9,999 permutations). Forest_nature4 denotes the four management categories: commercial, reserves, primeval-non-protected, and primeval-protected. Df, degrees of freedom; SS, sum of squares; R², proportion of variance explained by the grouping factor; F, pseudo-F ratio; p, permutation-based p value. PERMDISP tests whether within-group dispersion differs among categories; when it is significant, the PERMANOVA result can reflect differences in both group centroids and dispersion. Significance: *** p < 0.001, ** p < 0.01, * p < 0.05.

### Genetic diversity of study taxa

Population-level observed heterozygosity (Ho) ranged from 0.055 to 0.366 and expected heterozygosity (He) from 0.056 to 0.314. Species-level mean Ho spanned more than a threefold range, from 0.077 in *T. formicarius* to 0.290 in *E. ferrugineus*. The inbreeding coefficient (Fis) ranged widely among populations (−0.928 to 0.800), but species- and management-level means were close to zero or slightly negative (commercial: −0.021 ± 0.158; primeval-non-protected: −0.040 ± 0.157; primeval-protected: −0.021 ± 0.135; reserves: −0.003 ± 0.201), indicating a slight excess of heterozygotes rather than consistent inbreeding across most populations Supplementary Table S3.

#### Forest structural predictors of genetic diversity (H1)

The beta GLMM identified deadwood diversity as the only structural predictor with a significant within-species association with Ho (β = 0.041 ± 0.009, z = 4.44, 95% CI 0.0227–0.0586, p = 9.0 x 10⁻⁶; Table 3). Thus, populations in plots containing a more diverse mixture of deadwood categories tended to retain higher heterozygosity after accounting for species and site. Total deadwood volume, forest age, time since logging, old stump density, veteran tree count, and all pairwise management-category contrasts were non-significant (all p > 0.4). The Ho model had marginal R² = 0.0005 and conditional R² = 0.052, with no detected residual spatial autocorrelation (Moran’s I = 0.024, p = 0.352), indicating that the pooled association was statistically consistent but small. In the Gaussian Fis model, old stump density had a negative coefficient (β = −0.0152, SE = 0.0068, 95% CI −0.0286 to −0.0018, p = 0.026), whereas the other structural predictors and all management contrasts were non-significant. This Fis association remains provisional because diagnostics detected a small distributional deviation (KS p = 0.0495), significant outliers (p < 0.001), and conditional-quantile deviations, although dispersion (p = 0.678) and residual spatial autocorrelation (p = 0.454) were non-significant. Per-species regressions of Ho against deadwood diversity and time since logging are shown in Figures 4 and 5. Eleven of the 13 species had positive estimated slopes, but confidence intervals were broad. After Benjamini–Hochberg correction, only *B. reticulatus* retained a significant positive deadwood-diversity association (β = 0.027, adjusted p = 0.031), indicating that this species provided the clearest species-level support for the pooled pattern. *S. pectinicornis* showed the next-strongest positive estimate but remained uncertain after correction (β = 0.012, adjusted p = 0.079); estimates for pest species were small (all β < 0.005). Per-species relationships between Fis and deadwood diversity or time since logging, together with standardised effect-size plots for Ho and Fis, are presented in Supplementary Figures S2–S5.

**Figure 4.**
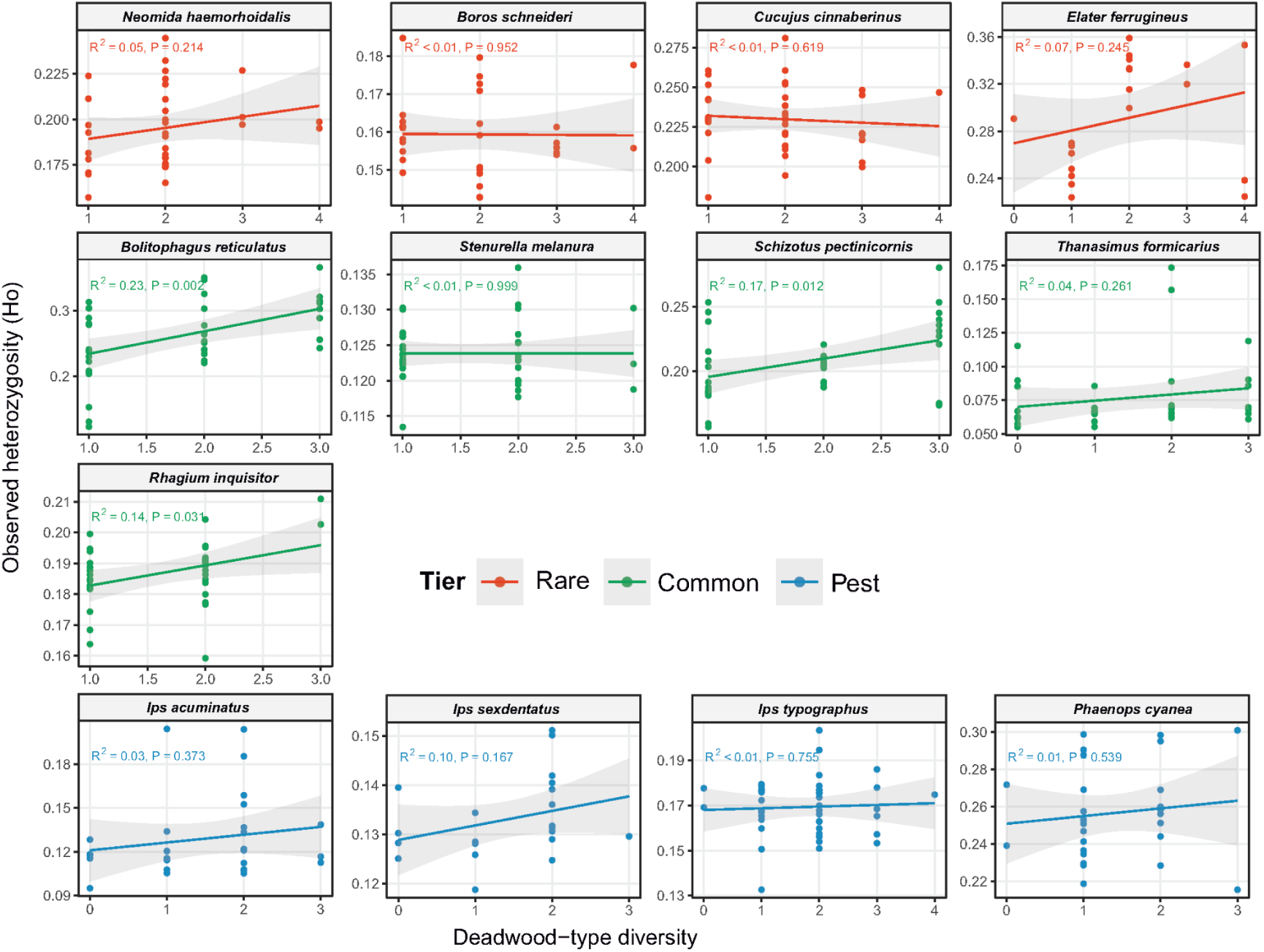
Per-species linear regressions of heterozygosity against deadwood diversity in examined saproxylic beetle species.

**Figure 5.**
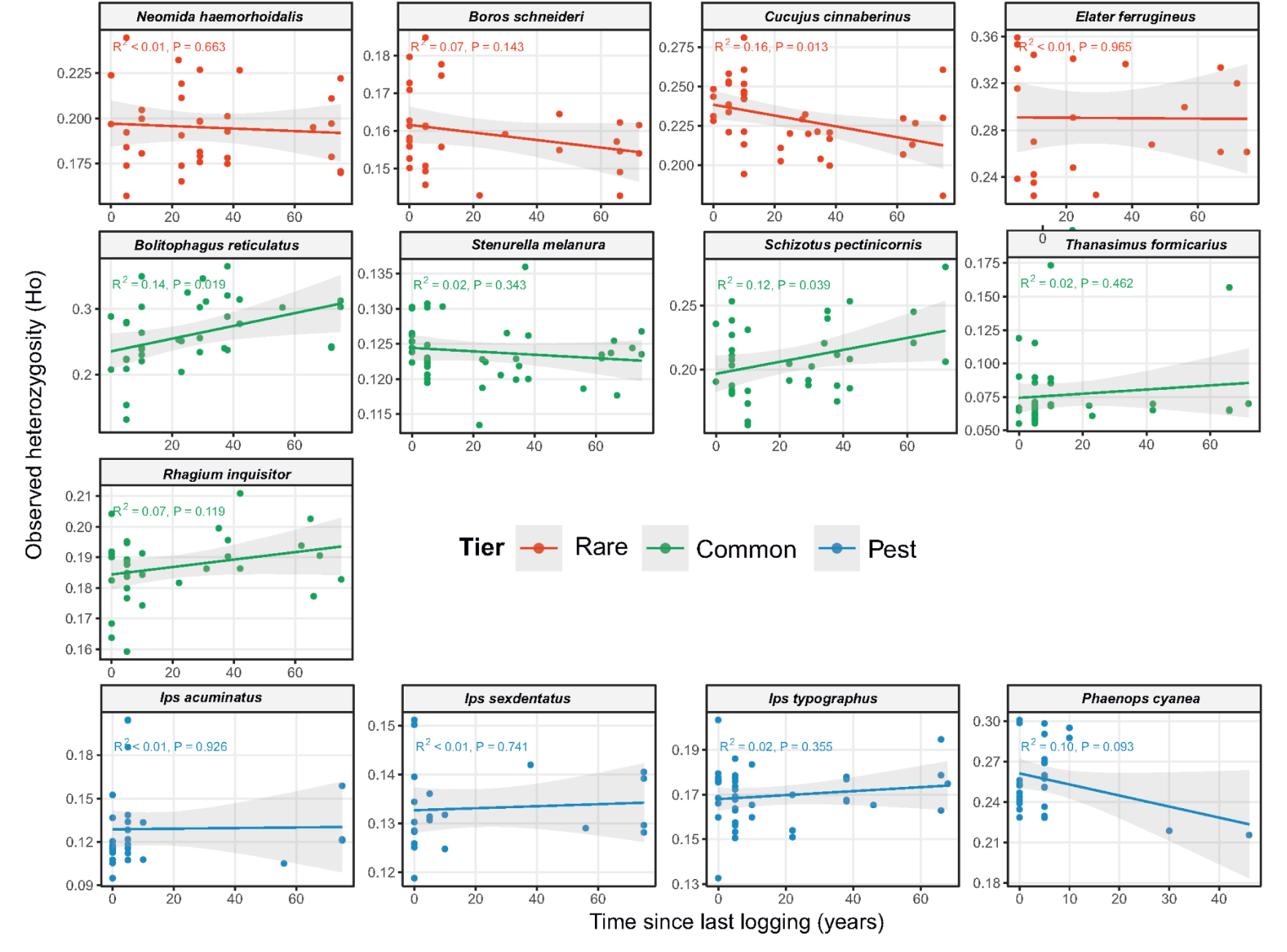
Per-species linear regressions of heterozygosity against time since latest logging in examined saproxylic beetle species.

**Table 3.** Beta Generalized Linear Mixed Models fixed-effects estimates. Predictors are z-scored. Tukey-adjusted pairwise contrasts on the management category shown below.

| Predictor | $\beta$ (logit) | SE | z | p |
| --- | --- | --- | --- | --- |
| Intercept | -1.544 | 0.119 | -13.02 | $< 2 \times 10^{-16} ***$ |
| V_total (z) | -0.0075 | 0.0092 | -0.82 | 0.414 |
| Max_age (z) | 0.0041 | 0.0102 | 0.40 | 0.691 |
| Latest_Logging (z) | 0.0079 | 0.0143 | 0.55 | 0.581 |
| Stumps_old (z) | 0.0069 | 0.0093 | 0.74 | 0.461 |
| Veterans (z) | -0.0001 | 0.0092 | -0.01 | 0.989 |
| Deadwood_diversity (z) | 0.041 | 0.009 | 4.44 | $9.0 \times 10^{-6} ***$ |
| Forest_nature4 (primeval-non-protected) | 0.037 | 0.024 | 1.54 | 0.124 |
| Forest_nature4 (primeval-protected) | 0.018 | 0.035 | 0.51 | 0.611 |
| Forest_nature4 (reserves) | 0.004 | 0.035 | 0.12 | 0.909 |
| Model fit |  |  |  |  |
| Species variance $\sigma^2$ | 0.179 | - | - | - |
| Dispersion (beta) | 239 | - | - | - |
| $R^2_m$ (marginal) | 0.0005 | - | - | - |
| $R^2_c$ (conditional) | 0.052 | - | - | - |
| Moran's I (residuals) | 0.024 | - | - | 0.35 |
| Contrast | Estimate | SE | z | p (Tukey) |
| commercial – primeval-non-protected | -0.037 | 0.024 | -1.54 | 0.415 |
| commercial – primeval-protected | -0.018 | 0.035 | -0.51 | 0.957 |
| commercial – reserves | -0.004 | 0.035 | -0.12 | 0.999 |
| primeval-non-protected – primeval-protected | 0.019 | 0.033 | 0.57 | 0.941 |
| primeval-non-protected – reserves | 0.033 | 0.034 | 0.97 | 0.766 |
| primeval-protected – reserves | 0.014 | 0.026 | 0.53 | 0.951 |

#### Effective migration surfaces among ecological strategies (H2)

Estimated effective migration surfaces revealed highly species-specific patterns, rejecting a uniform community-wide response to landscape fragmentation (Figure 6). The 13 species clustered into three primary macro- geographic migration scenarios. First, regional lineage barriers were identified: a pronounced latitudinal migration barrier through central Poland occurred in *C. cinnaberinus*, *N. haemorrhoidalis*, and *P. cyanea*, isolating western populations from eastern genetic reservoirs. *T. formicarius* displayed an alternative pattern, with a broad high-migration corridor across the southern mountain range bounded by a pronounced barrier in the northern lowlands. Second, asymmetrical corridors were found: *I. acuminatus* showed an inverted pattern, in which the western region functioned as a deep gene-flow barrier while the eastern and northeastern lowlands formed a broad migration corridor. *B. schneideri* and *I. typographus* exhibited diffuse high-migration corridors across the central and northeastern lowlands, indicating high structural connectivity across unfragmented habitats. Third, *B. reticulatus* and *S. melanura* displayed mosaic-like patches of alternating corridors and barriers, with *S. melanura* tracking a distinct barrier along the southern montane border, whereas *E. ferrugineus* and *I. sexdentatus* exhibited featureless migration surfaces closely conforming to standard isolation-by-distance expectations.

**Figure 6.**
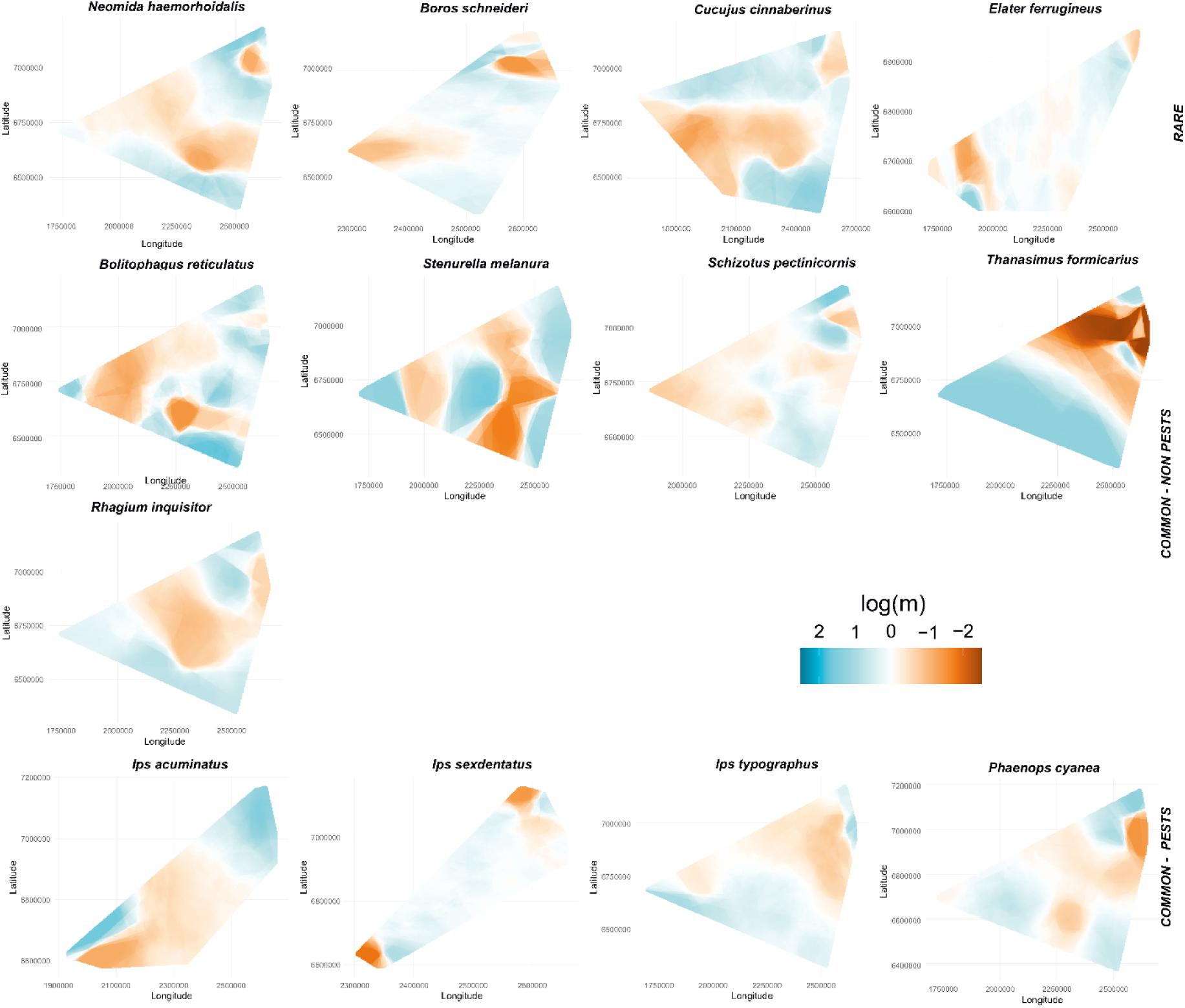
Estimated effective migration surfaces (EEMS) for 13 saproxylic beetle species. Maps display spatial deviations from isolation-by-distance expectations across Central Europe. Blue shading denotes corridors of elevated gene flow; orange shading identifies barriers to migration. Teal/blue (log(m) > 0, corridors) through white (log(m) = 0) to orange (log(m) < 0, barriers).

#### Environmental-response congruence across ecological tiers (H3)

Tier comparisons showed the highest mean Ho in rare relict species, intermediate values in common forest taxa, and the lowest values in pest species (Figure 3; all pairwise Wilcoxon tests p < 10⁻⁵ after Benjamini–Hochberg adjustment; Supplementary Table S1). Fis showed the inverse pattern. The four management categories did not differ significantly in either Ho or Fis after Tukey adjustment (all pairwise contrasts p > 0.4; Figure 3; Table 3). In the rare–common model, five of six predictor x tier interactions were non-significant, indicating no detectable tier difference in the slopes for total deadwood volume, forest age, old stump density, veteran tree density, or deadwood diversity (Table 4). Time since logging was the sole exception (χ² = 4.56, p = 0.033): the estimated slope was positive for common species (β = 0.033, 95% CI −0.006–0.072) and negative for rare species (β = −0.016, 95% CI −0.054–0.022), although both tier-specific confidence intervals included zero. Sensitivity analyses using two alternative binary management codings produced the same conclusion for time since logging (χ² = 4.56–4.61, p = 0.032–0.038). The deadwood diversity x tier interaction remained borderline (p = 0.049–0.053), so it is interpreted as uncertain rather than as clear evidence of tier divergence.

**Table 4.**
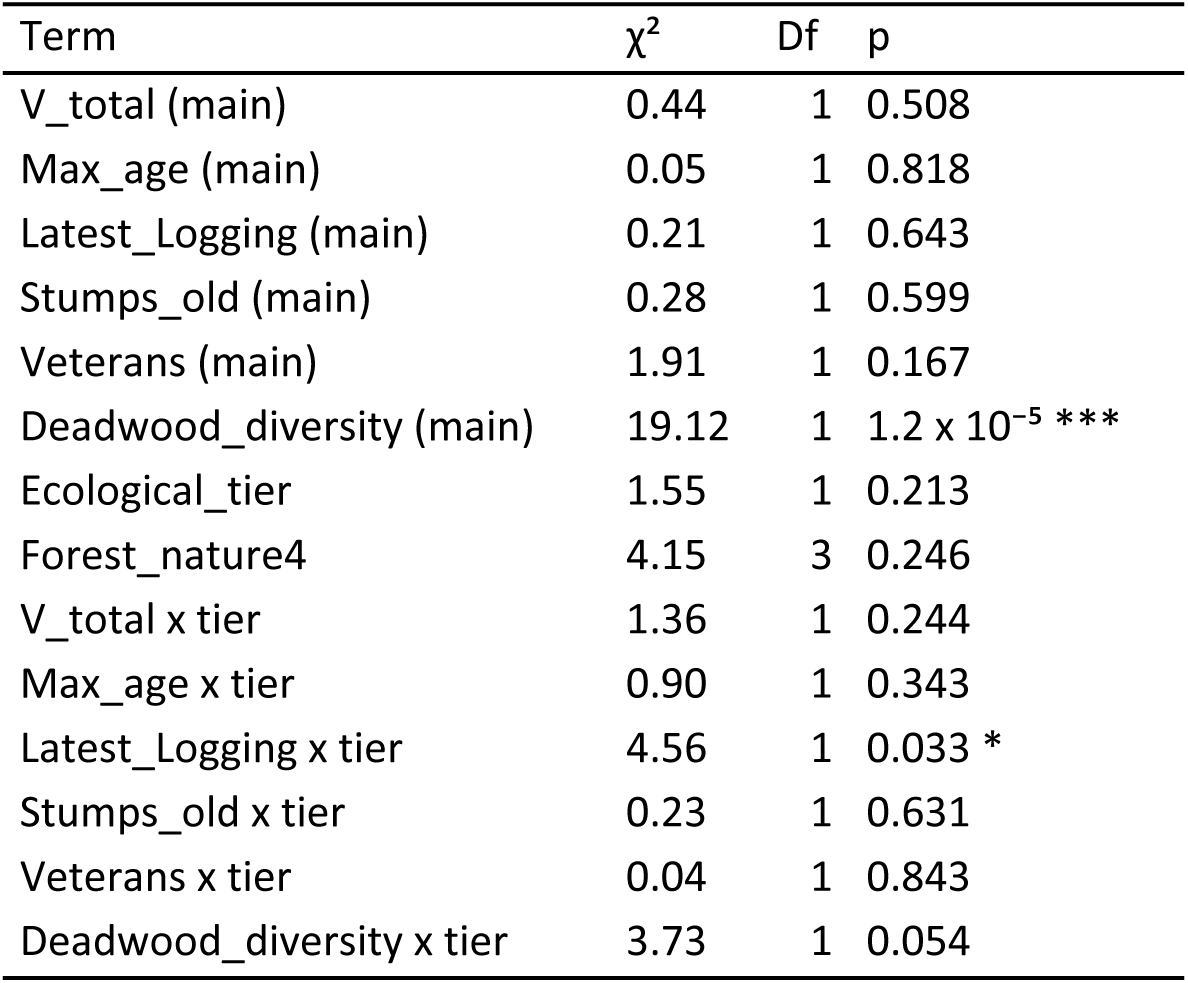
Type-II Wald χ² tests for predictor x ecological-tier interactions in the rare–common mixed model (N = 312 site-level population records from 9 species). Non-significant interactions indicate no detectable difference between tier-specific slopes; the significant time-since-logging interaction indicates a difference between tiers.

## Discussion

The genetic diversity of saproxylic beetle assemblages is shaped by processes operating at multiple spatial scales, although only a subset of the forest characteristics examined were associated with genetic variation. At the local (plot) scale, observed heterozygosity increased with deadwood diversity but not with total deadwood volume or other structural predictors, identifying deadwood diversity as the principal habitat attribute associated with within-population genetic diversity. At broader spatial scales, macro-geographic landscape configuration influenced gene flow in a strongly species-specific manner, indicating that no single connectivity pattern characterises the entire assemblage. These genetic responses were further mediated by species-specific traits related to their commonness, dispersal ability, and sensitivity to anthropogenic forest modification. Although comparisons between managed and old-growth forests suggested that long-term forest protection can enhance the genetic diversity of rare and specialist saproxylic beetles, management-category effects on observed heterozygosity and inbreeding were not significant after correction for multiple testing and were not consistent across the assemblage. We therefore discuss the local- and regional-scale drivers separately before considering their implications for the conservation of saproxylic beetles and the maintenance of their genetic diversity.

### Deadwood diversity and within-population genetic variation

Across 13 saproxylic beetle species and 430 sampling sites, the beta GLMM identified deadwood diversity as the only structural predictor with a consistent within-species association with observed heterozygosity. Total deadwood volume was not significant. This difference between deadwood amount and diversity is ecologically meaningful: a forest containing a relatively large quantity of homogeneous deadwood may provide less microhabitat heterogeneity than a forest with less deadwood distributed across multiple tree species, decay classes, and physical configurations (Müller & Bütler 2010; Bouget et al. 2014). Saproxylic communities depend on this heterogeneity because species specialise on different decay stages and substrate types (Stokland et al. 2012; Bauhus et al. 2018). The genetic result therefore extends a well-established community-level pattern: structural variety, rather than bulk biomass alone, supports saproxylic genetic diversity.

Neither heterozygosity nor inbreeding differed significantly among the four management categories after Tukey adjustment. Coarse management categories may therefore capture less within-species genetic variation than direct measures of local structural quality (Lowe & Allendorf 2010). This complements the significant management effect detected at the between-population fixation index (FST) level by Amer et al. (2026): between-population differentiation can respond to management-driven fragmentation, whereas within-population heterozygosity can respond to fine-scale structural quality. This scale-dependent contrast is consistent with landscape genetics, in which FST reflects connectivity and heterozygosity reflects local habitat quality (Manel & Holderegger 2013; Hohenlohe et al. 2021).

Multivariate ordination showed that diversity (heterozygosity) and inbreeding loaded on different axes (PC1 and PC2), indicating that heterozygosity loss and within-population inbreeding are at least partly decoupled. A population can lose heterozygosity through drift while retaining random mating, or accumulate inbreeding while heterozygosity remains buffered by past gene flow (Crow & Kimura 2010). Treating these indices as complementary rather than redundant measures of genetic condition is therefore appropriate for monitoring saproxylic beetles.

### Life-history traits and macro-geographic resistance

Effective migration surfaces varied among species, consistent with differences in life history and in the distribution of suitable forest habitat (Chambers et al. 2025; Kobayashi & Sota 2019). Pest taxa also showed broad geographic corridors and barriers. The central and western barriers estimated for *P. cyanea* and *I. acuminatus* indicate restricted regional gene flow despite their high local abundance. One possible explanation is that the distribution and management history of host trees, including rotation patterns in monocultures, divide otherwise abundant populations (Lagisz et al. 2010; Zytynska et al. 2018).

Outbreak dynamics may also influence genetic structure in pest species. Although several of these taxa show low differentiation across their European or Central European ranges (Krovi et al. 2025; Amer et al. 2026), outbreaks can produce temporary local structure (Papek et al. 2024). In the species-level analyses, heterozygosity in *P. cyanea* increased with veteran tree density. Because *P. cyanea* uses stressed pine, older pine stands may provide more persistent host resources than short-lived outbreak patches; this interpretation remains tentative because it is based on a single-species association.

Rare old-growth specialists did not share a single regional pattern. *C. cinnaberinus* and *N. haemorrhoidalis* showed regional splits between western populations and eastern old-growth areas, whereas the *E. ferrugineus* surface was close to the isolation-by-distance expectation. The former patterns may include historical structure: *C. cinnaberinus* comprises two mitochondrial lineages in mainland Europe (Sikora et al. 2023). The weak EEMS structure in *E. ferrugineus* is less readily explained by its association with the low-dispersal hermit beetle *Osmoderma barnabita*, although *E. ferrugineus* also preys on other, more widespread Cetoniinae (Oleksa et al. 2013; Zauli et al. 2014; Chiari et al. 2013). The strong genetic isolation of the Holy Cross Mountains population of *B. schneideri* is consistent with the findings of Amer et al. (2026) and with the distinct genetic distinctiveness reported for another boreal relict beetle, *Cucujus haematodes* Erichson, 1845 (Kadej et al. 2022). Together, these findings highlight the exceptional conservation value of these primeval forests.

Among species with featureless or mosaic-like surfaces, *B. reticulatus*, *S. melanura*, and *I. sexdentatus* deserve brief mention. *B. reticulatus* is a generalist fungivore on tinder fungus (*Fomes fomentarius* (L.) Fr.), which occurs on several host tree species across Europe. Its mosaic may reflect the local distribution of fungal substrate, consistent with the species’ phylogeography (Knutsen et al. 2000; Eberle et al. 2021). Adult *S. melanura* use floral resources, so its southern barrier may be related to discontinuities in Carpathian montane meadows or to recently described mitochondrial haplogroups rather than to forest structure alone (Melnyk et al. 2026). *I. sexdentatus* has been reported as panmictic at the European scale (Avtzis et al. 2019). Its Polish surface lacked strong barriers or corridors; here, “near-IBD” refers to that near-null migration surface rather than to evidence for pronounced isolation by distance.

These migration patterns are partly concordant with the phylogeography of the studied species in Central Europe: relict and stenotopic species tend to show strong population distinctiveness, whereas generalists, including pests, have widespread genetic lineages (Kajtoch et al. 2022; Krovi et al. 2025). Contemporary spatial genetic structure in saproxylic beetles therefore reflects both historical and recent habitat change.

Mean heterozygosity was highest in rare relict species, intermediate in common species, and lowest in pest species (Figure 3), contrary to the expectation of reduced diversity in rare specialists. The four rare species are habitat specialists but remain broadly distributed in Europe, and the sampled primeval forest remnants may retain old genetic variation. Sampling may also contribute: rare specialists were usually collected in high-quality refugia, whereas common species were sampled across a wider range of habitat conditions. For *Ips spp.*, recurrent outbreaks can reduce effective population size even when census abundance is temporarily high (Bertheau et al. 2013; Avtzis et al. 2019). These explanations are not mutually exclusive, and the tier comparison should not be interpreted as a general property of all rare, common, or pest saproxylic beetles.

Five of the six predictor x tier tests found no detectable difference between rare and common species. Time since logging was the exception: heterozygosity tended to increase with longer logging-free intervals in common species, whereas the slope for rare relicts was close to zero and uncertain. Rare species in this dataset already occurred mainly in old, relatively stable refugia, including old-growth remnants within commercial forests, which may limit the additional association with time since logging (Oleksa et al. 2015).

The time-since-logging result suggests that longer rotations and post-harvest recovery may be more relevant to common species than to the sampled rare relicts. For the other structural variables, the analyses did not detect different slopes between tiers. Increasing deadwood diversity may thus benefit both groups, although the species-level estimates were imprecise. Positive associations between veteran tree density and inbreeding in some rare specialists should not be read as evidence against veteran tree retention; they may instead reflect persistent use of localised microhabitats by low-dispersal species (Oleksa et al. 2015; Horák 2017).

### Limitations

Several species-level results require caution. *T. formicarius* included populations with very high positive Fis, which may reflect null alleles, mixing of differentiated samples, or another technical source of heterozygote deficit. The *I. typographus* EEMS surface and *T. formicarius* heterozygosity estimates are therefore treated as low confidence pending targeted re-sequencing. EEMS also estimates time-integrated effective migration and cannot separate current fragmentation from older biogeographic structure (Petkova et al. 2016; Bilgin 2011). The central-Poland barriers in *C. cinnaberinus* and *N. haemorrhoidalis* may thus include a signal of post-glacial history. Distinguishing contemporary and historical effects would require demographic-history analyses that were not part of this study. Finally, per-species sample sizes ranged from 21 to 42 populations. The pooled models use information across species, but most individual species tests had low precision; only the *B. reticulatus* association remained significant after correction for multiple testing.

### Conservation Implications

These implications apply to protected rare species associated with primeval forests and to common species that depend on old-growth structures. Pest species are not conservation targets, although their EEMS surfaces provide information about host distribution and management history. Deadwood diversity was associated with heterozygosity, whereas total deadwood volume was not. Existing volume guidelines remain relevant to saproxylic assemblages (Müller & Bütler 2010; Siitonen & Saaristo 2000; Lassauce et al. 2011; Thorn et al. 2020), but our results suggest that managers should also retain various tree species, decay classes, and physical forms of deadwood (e.g. snags and logs). Examples from the field associations include lying decayed logs for *C. cinnaberinus* and *S. pectinicornis*; veteran deciduous trees and hollows for *E. ferrugineus*; deciduous trees bearing tinder fungus (*Fomes fomentarius*) for *N. haemorrhoidalis*; and standing coniferous deadwood for *B. schneideri*.

The interaction with time since logging was supported statistically, but the tier-specific slopes were individually uncertain. Nevertheless, the positive estimated slope for common species suggests that longer rotations and post-harvest recovery may benefit this group (Table 4). For rare relicts already concentrated in stable old-growth refugia, maintaining the refugia is likely to be the most important, although ceasing logging between refugia may be beneficial for restoring their connectivity. Although our analyses do not test reserve size or configuration directly, they provide relevant context for the SLOSS debate (Diamond 1975; Fahrig et al. 2022) by showing that spatial genetic patterns differ among species. These findings are consistent with a complementary conservation strategy that protects large, contiguous primeval forest complexes—such as Białowieża Forest, the Carpathians, and the Holy Cross Mountains—while maintaining smaller, structurally diverse habitat patches that may contribute to species-specific connectivity. Because EEMS describes spatial variation in effective migration rather than the relative performance of alternative reserve designs (Petkova et al. 2016), the effectiveness of these complementary elements requires direct evaluation.

In commercial forests, networks of patches containing veteran trees, diverse deadwood, and species-specific substrates could complement protected refugia. Similarity between populations in the Carpathians and the Holy Cross Mountains may reflect historical continuity of old-growth conditions, potentially through the Roztocze Upland (Papis & Mokrzyski 2015), but this proposed connection requires direct testing.

The association with logging history supports minimising disturbance in protected areas and allowing recovery after harvest. The EU Biodiversity Strategy for 2030 (European Commission 2020) calls for all remaining primeval and old-growth forests to be included in a strictly protected network. Given the species-specific nature of landscape resistance revealed by our EEMS models, effective multi-species conservation will require spatially resolved genomic monitoring to identify and prioritise corridor restoration across the Central European forest network. Until now, such genetic-monitoring evidence has been available mostly for charismatic vertebrates, such as large carnivores; this study extends the same principle to invertebrate keystones of European forests.

## Acknowledgements

This research was funded by the National Science Centre, Poland (NCN grant UMO-2021/43/B/NZ9/00991). The authors are sincerely grateful to the Polish nature protection authorities for granting permission to conduct fieldwork in protected forest complexes and to collect protected species.

## Ethical Statement

Collection of protected beetle species and in protected areas was executed according to permissions granted by the Polish Ministry of Environment (DOP-1.61.61.2021.TP 1516071.4994962.3977511 and DOP-1.61.61.2021.TP 1516071.4994586.3977515), the General Directorate of Environmental Protection (DZP-WF.6401.68.2021.AS) and the Regional Directorates of Environmental Protection in Białystok (WPN.6205.74.2021.MM), Rzeszów (WPN.6205.116.2021.MP.2 and WPN.6205.114.2021.ŁL.2), Kielce (WPN.I.6205.1.49.2021.EJP.2) and Wrocław (WPN.6205.143.2021.AR).

## Data Availability Statement

Raw sequencing data (FASTQ files) and associated sample metadata have been deposited in the NCBI Sequence Read Archive (SRA) under BioProject PRJNA1399704. Filtered single-nucleotide-polymorphism datasets generated from ddRAD-seq for all saproxylic beetle species are available as species-specific Variant Call Format (VCF) files in RepoOD of the Institute of Systematics and Animal Evolution PAS (https://doi.org/10.18150/MYUQJG). All forest and ecological field measurements (CSV) are submitted as supplementary data along with the manuscript. Analysis code is archived on GitHub (https://github.com/SarvaRam/saproxylic-genetics-poland) under the MIT licence; a Zenodo DOI will be minted at v1.0.0.

## Author Contributions

R.S.K.: Conceptualisation, Sampling, Laboratory resources, Formal analysis, Investigation, Writing - original draft. N.R.A.: Formal analysis, review & editing. A.W., M.O., T.Sz.: Laboratory resources, review & editing. R.P., M.K., T.J., A.S.: Sampling, Investigation, review & editing. Ł.K, M.O. Conceptualisation, Supervision, Funding acquisition, Writing - review & editing.

### Competing Interests

The authors declare no competing interests.

Article Impact Statement Deadwood diversity drives beetle genetic diversity, but species-specific migration profiles require scale- sensitive conservation planning.

## Supplementary Tables and Figures

**Supplementary Figure S1.**
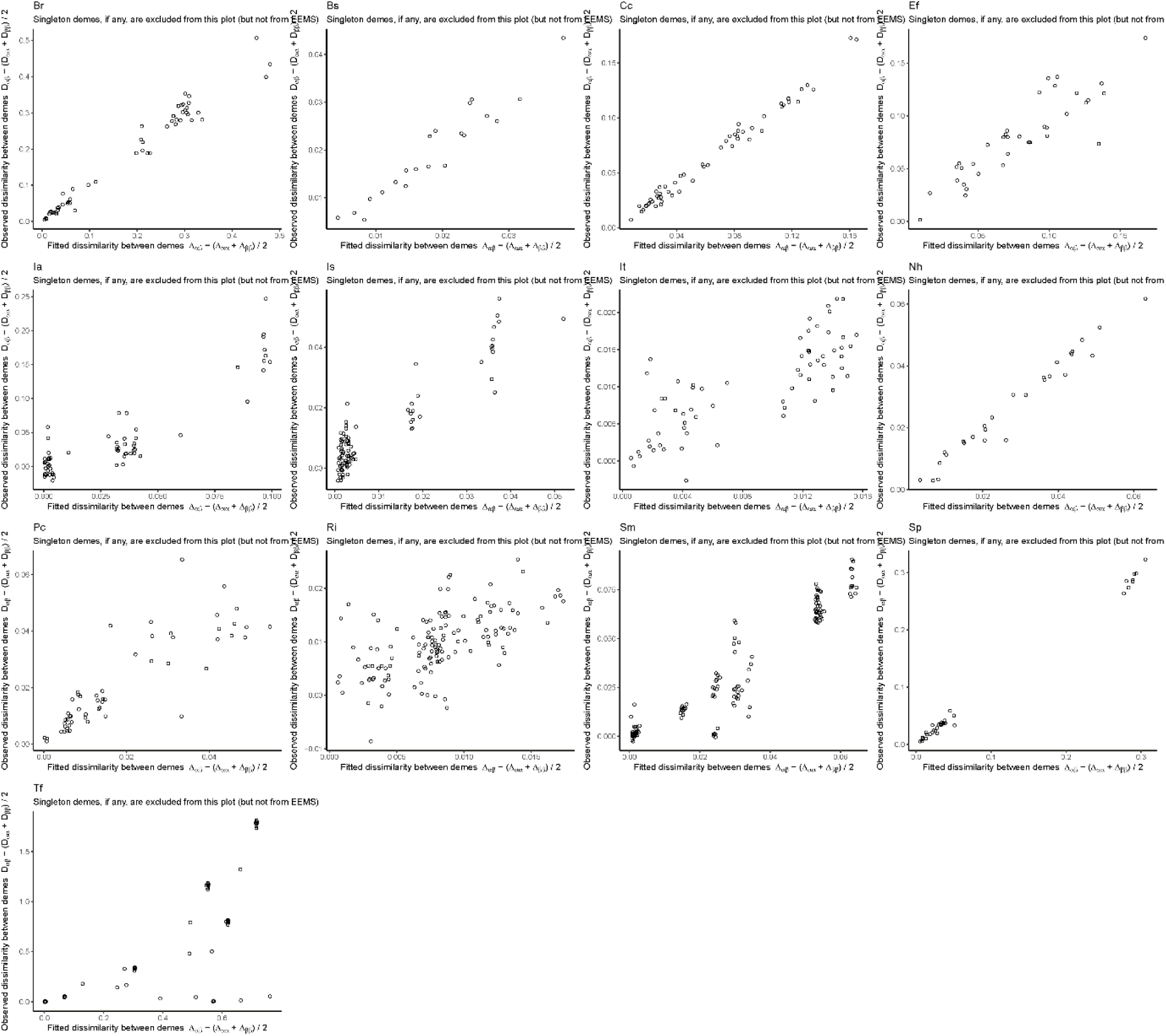
Observed versus fitted between-deme genetic dissimilarities for the 13 estimated effective migration surface (EEMS) models. Each panel represents one species, and each point represents a pair of sampled demes. Singleton demes, where present, were excluded from these diagnostic plots but retained in the EEMS analyses.

**Supplementary Figure S2.**
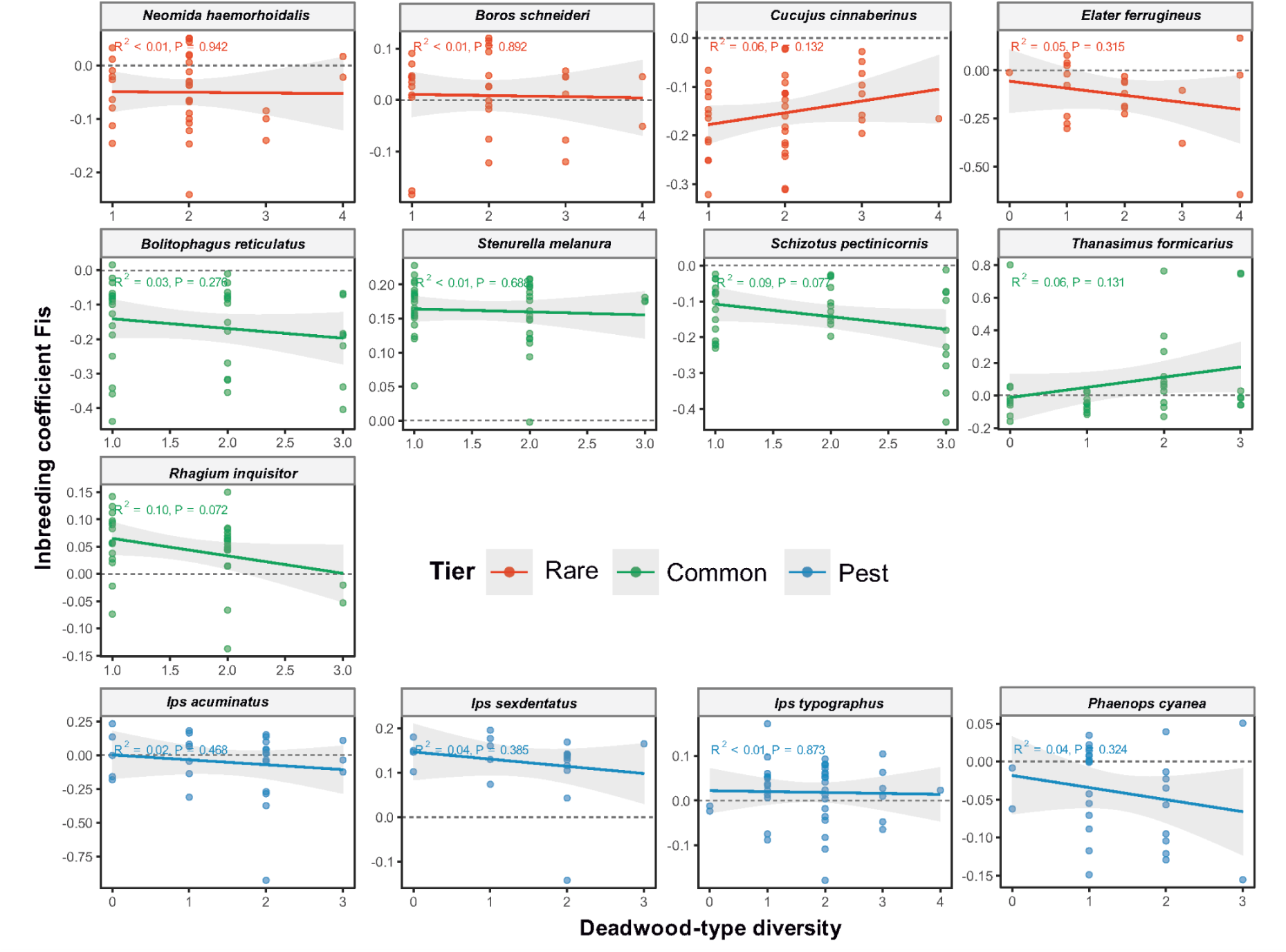
Per-species relationships between the inbreeding coefficient (Fis) and deadwood diversity. Points represent sampled populations; solid lines and grey bands show ordinary least-squares fits and 95% confidence intervals, respectively. The horizontal dashed line marks Fis = 0. Species and fitted lines are coloured by ecological tier: rare taxa in orange, common taxa in green, and pest taxa in blue. Panel annotations report unadjusted R² and p values.

**Supplementary Figure S3.**
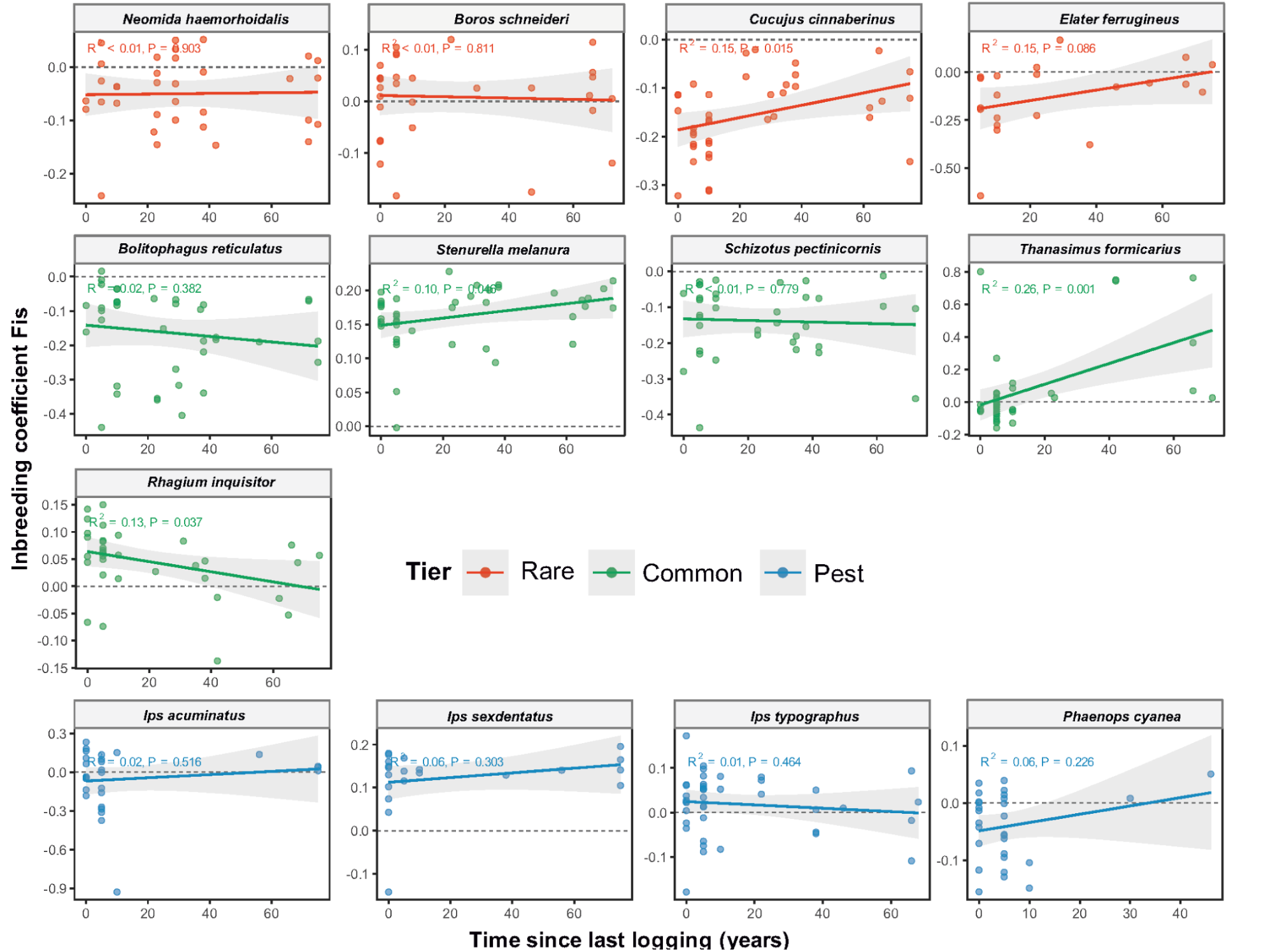
Per-species relationships between the inbreeding coefficient (Fis) and time since the most recent logging event. Points represent sampled populations; solid lines and grey bands show ordinary least-squares fits and 95% confidence intervals, respectively. The horizontal dashed line marks Fis = 0. Species and fitted lines are coloured by ecological tier: rare taxa in orange, common taxa in green, and pest taxa in blue. Panel annotations report unadjusted R² and p values.

**Supplementary Figure S4.**
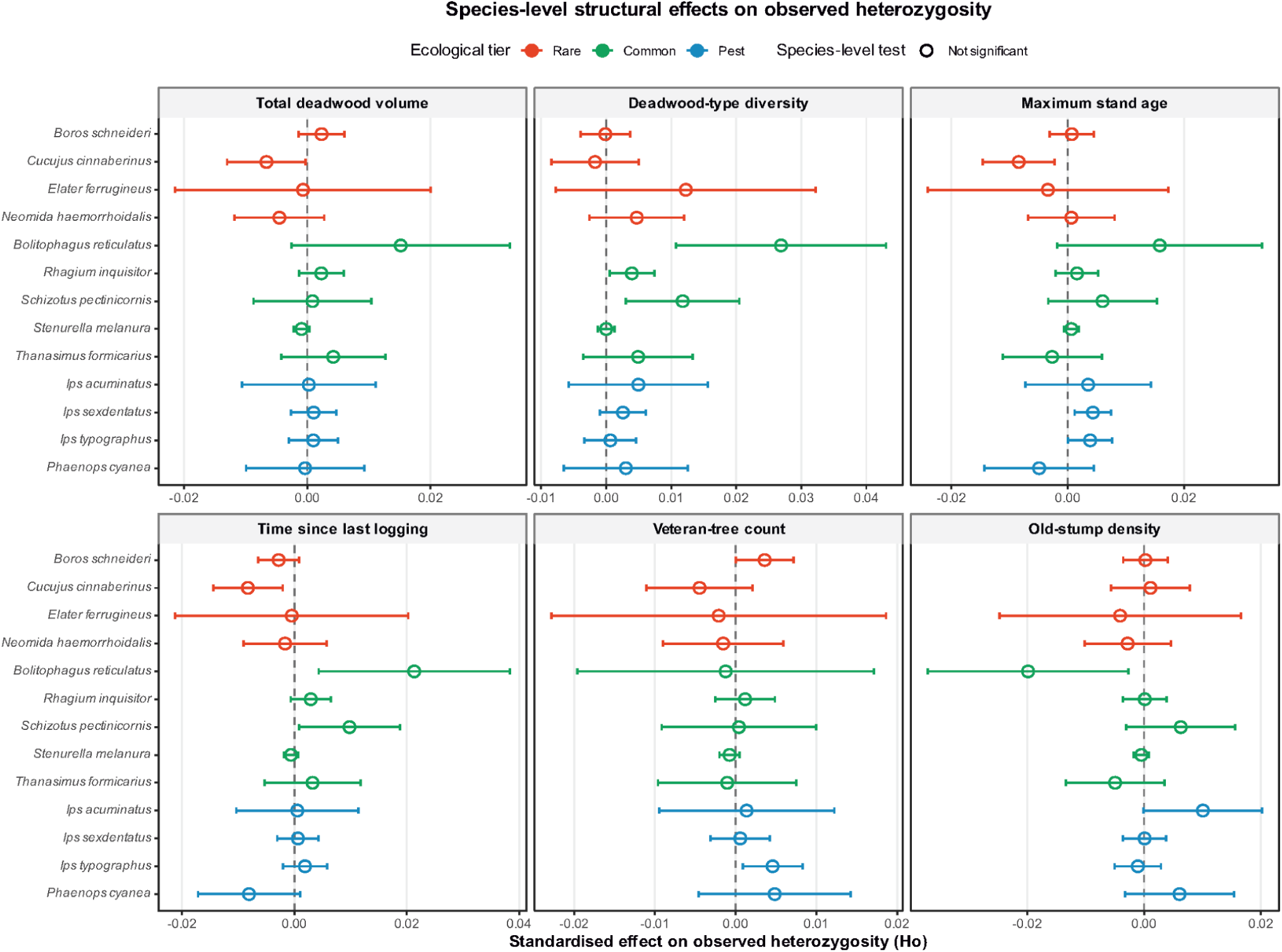
Standardised species-level effects of six forest structural predictors on observed heterozygosity (Ho). Circles show slope estimates and horizontal bars show 95% confidence intervals; the vertical dashed line marks zero effect. Estimates are coloured by ecological tier: rare taxa in orange, common taxa in green, and pest taxa in blue. Open circles denote non-significant species-level tests.

**Supplementary Figure S5.**
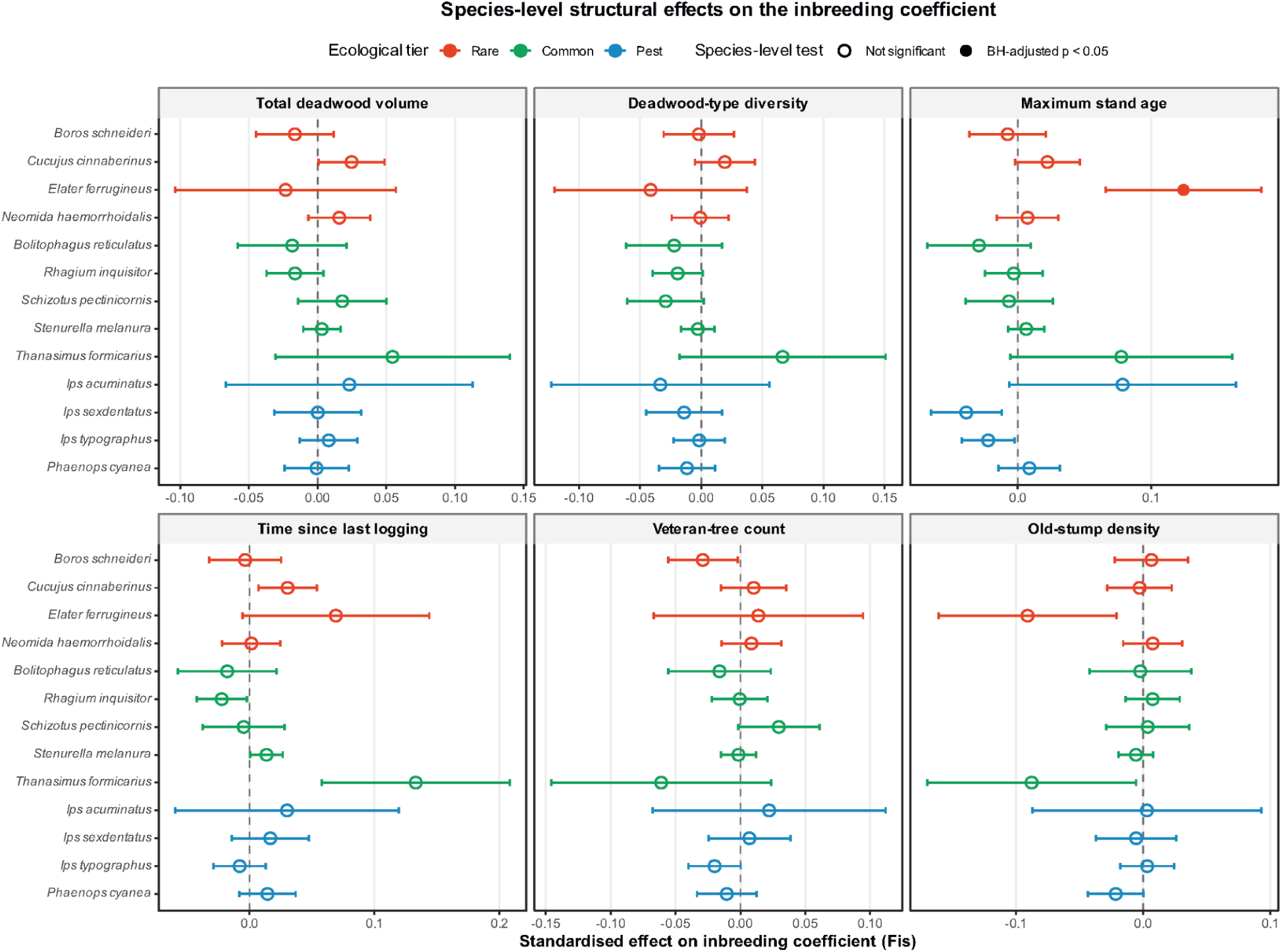
Standardised species-level effects of six forest structural predictors on the inbreeding coefficient (Fis). Circles show slope estimates and horizontal bars show 95% confidence intervals; the vertical dashed line marks zero effect. Estimates are coloured by ecological tier: rare taxa in orange, common taxa in green, and pest taxa in blue. Open circles denote non-significant tests, whereas filled circles denote Benjamini-Hochberg-adjusted p < 0.05 across the full 78-test species x predictor family.

**Table S1.**
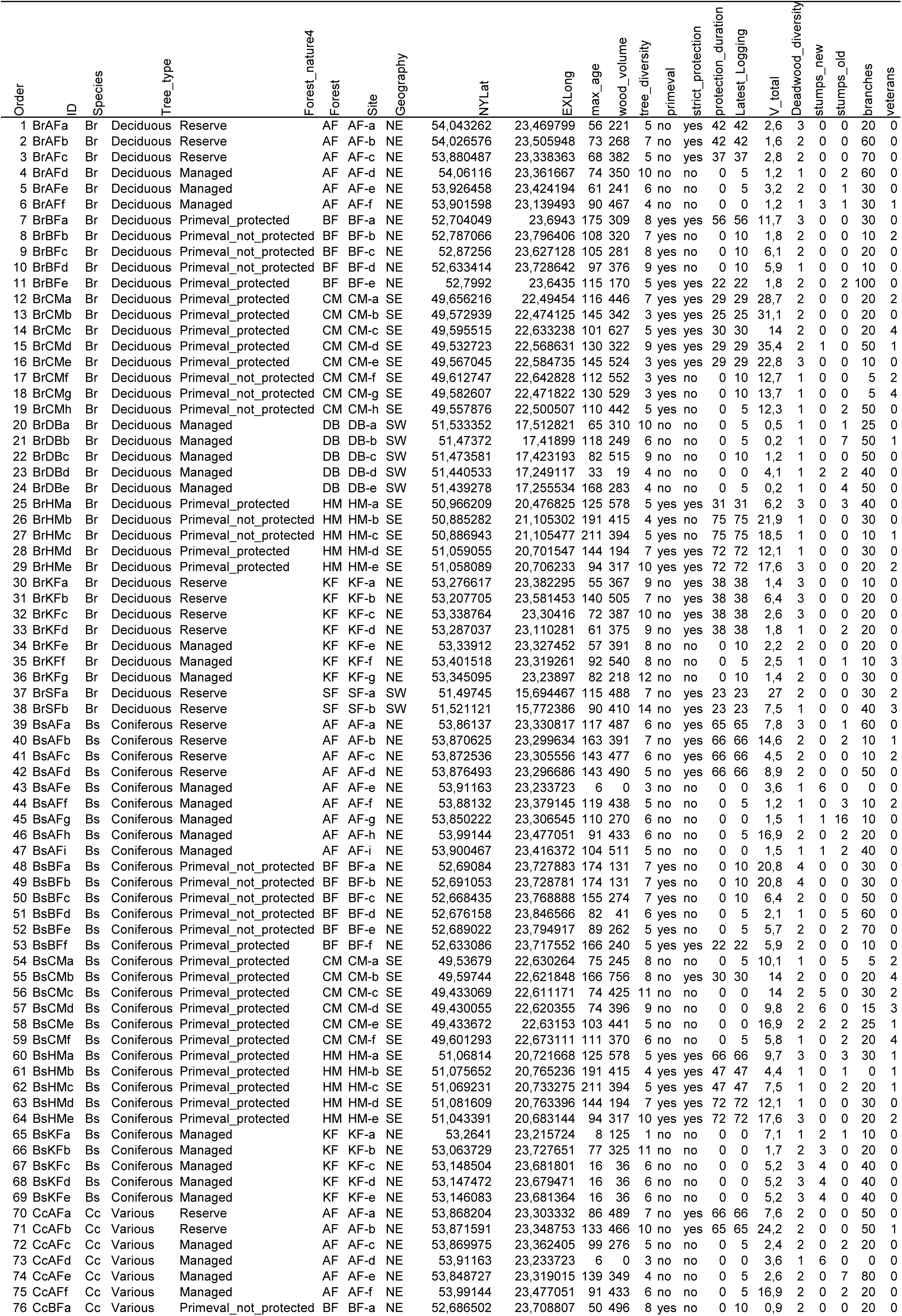
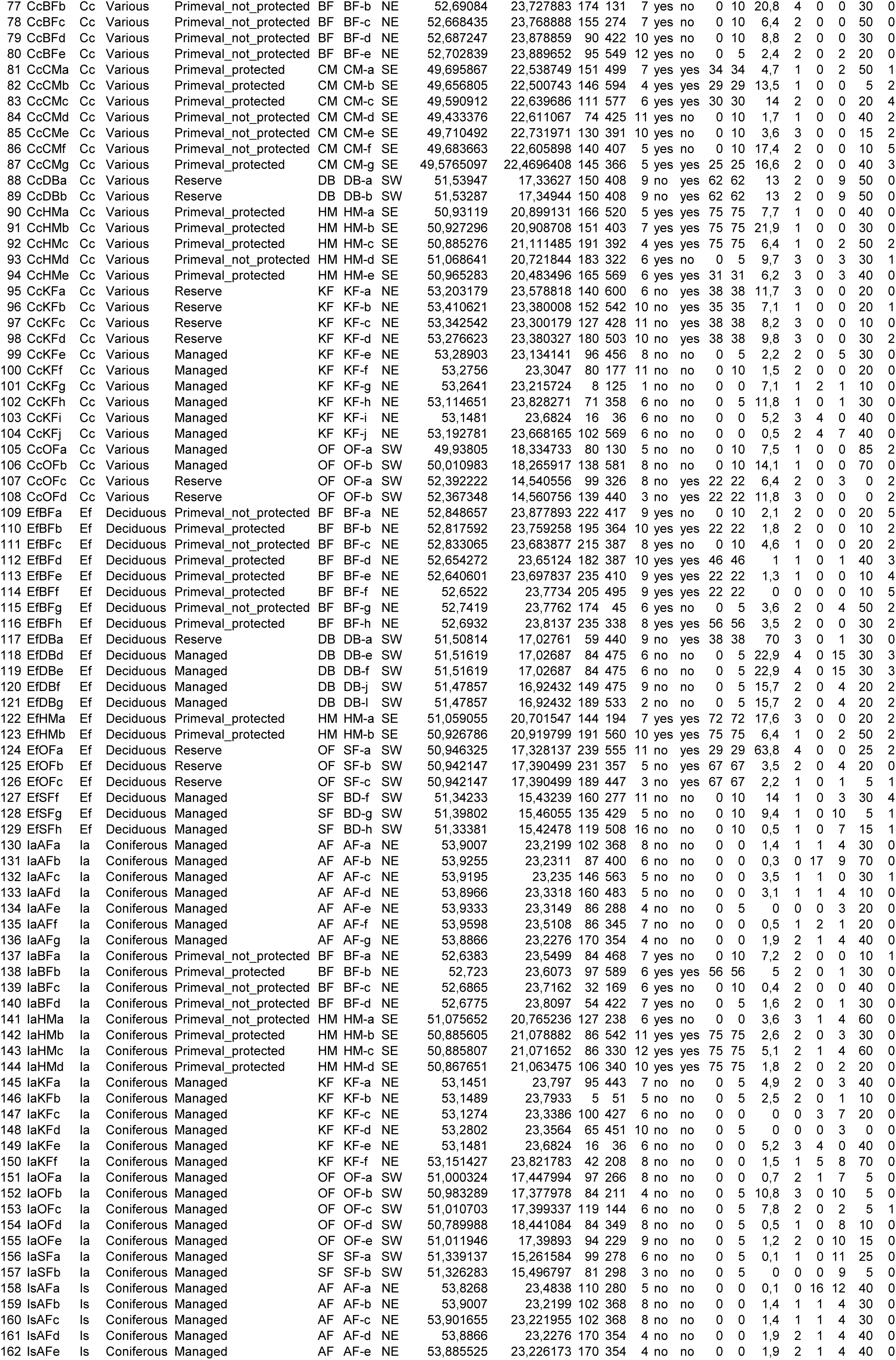
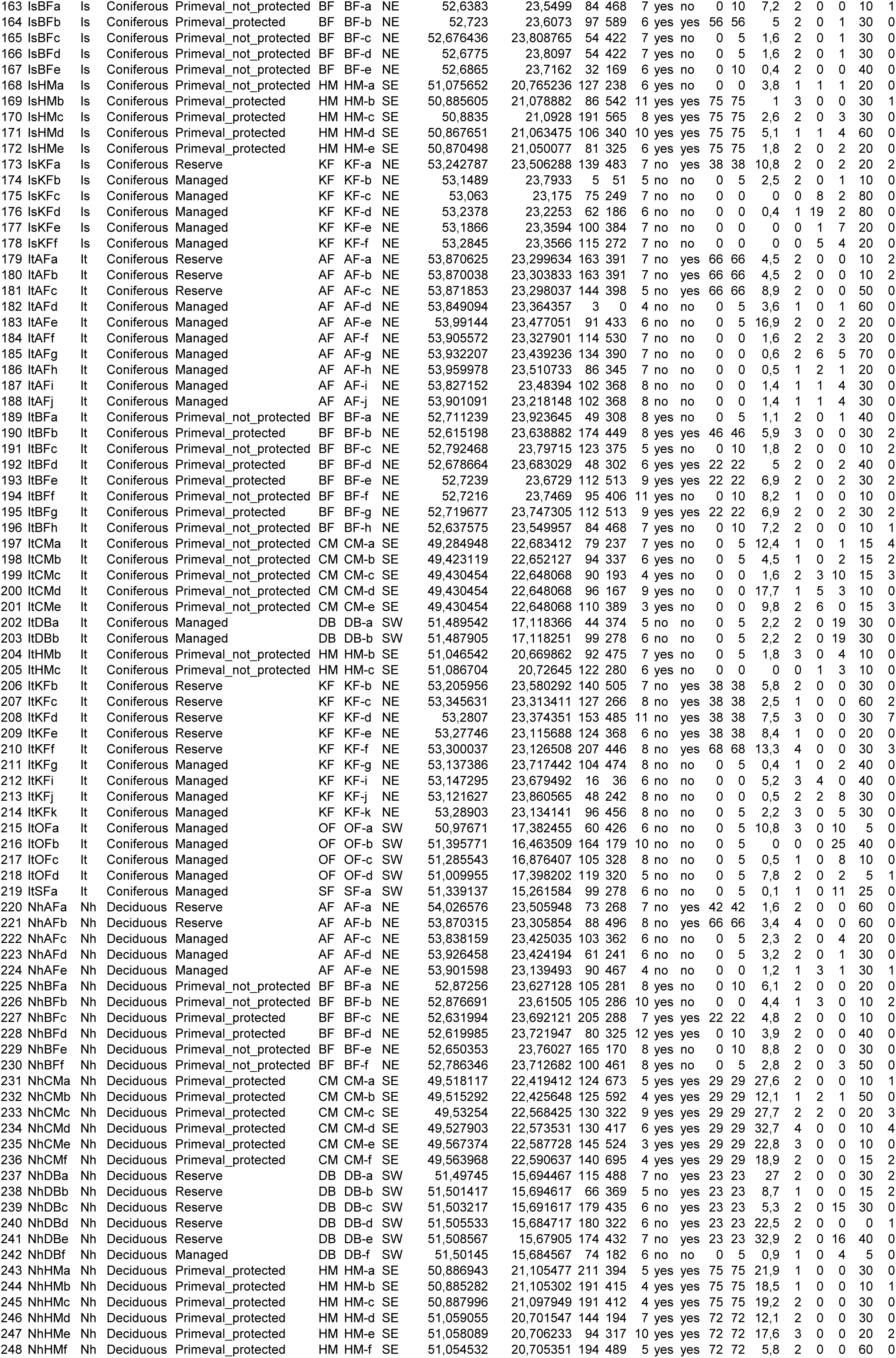
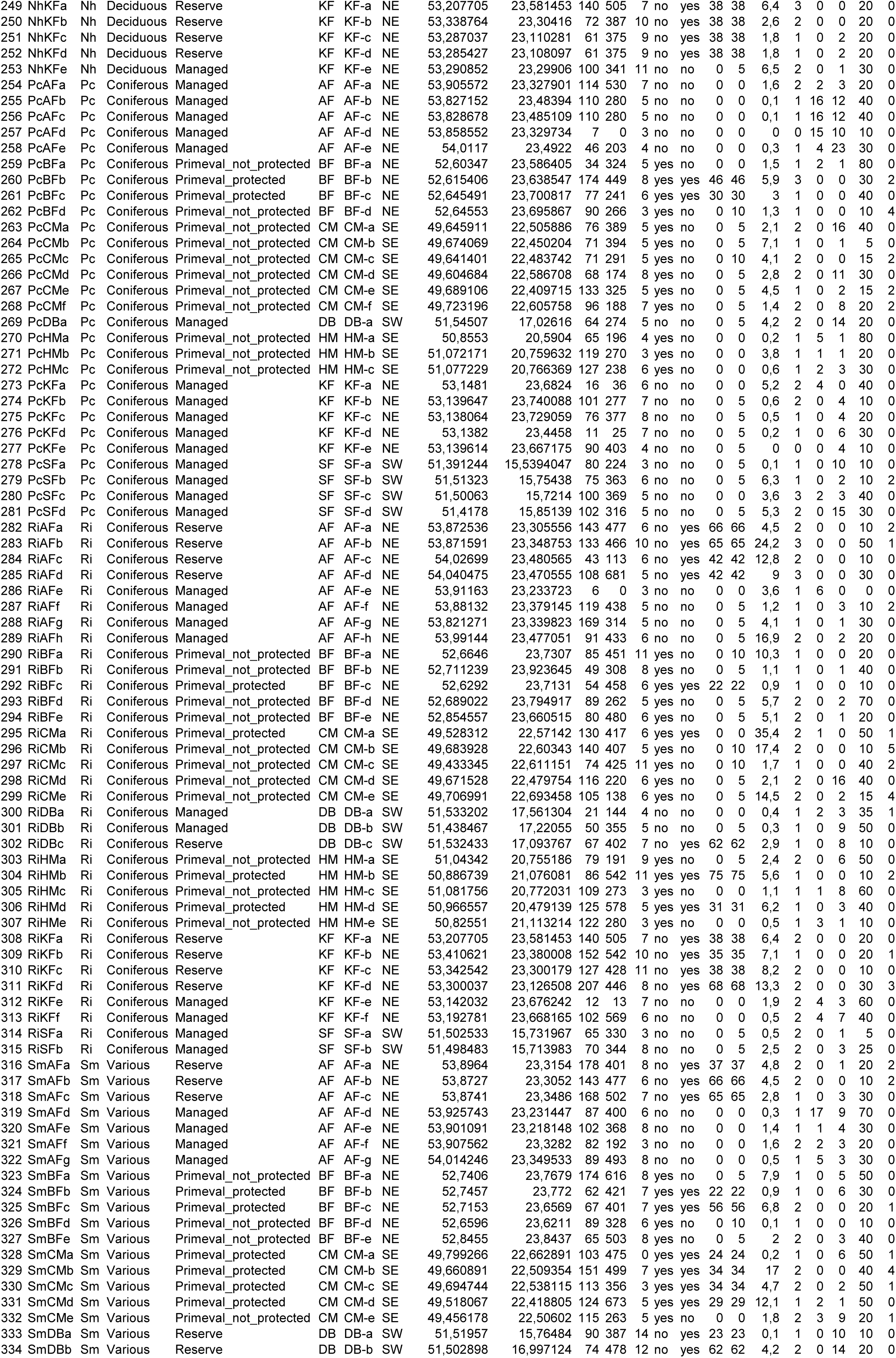
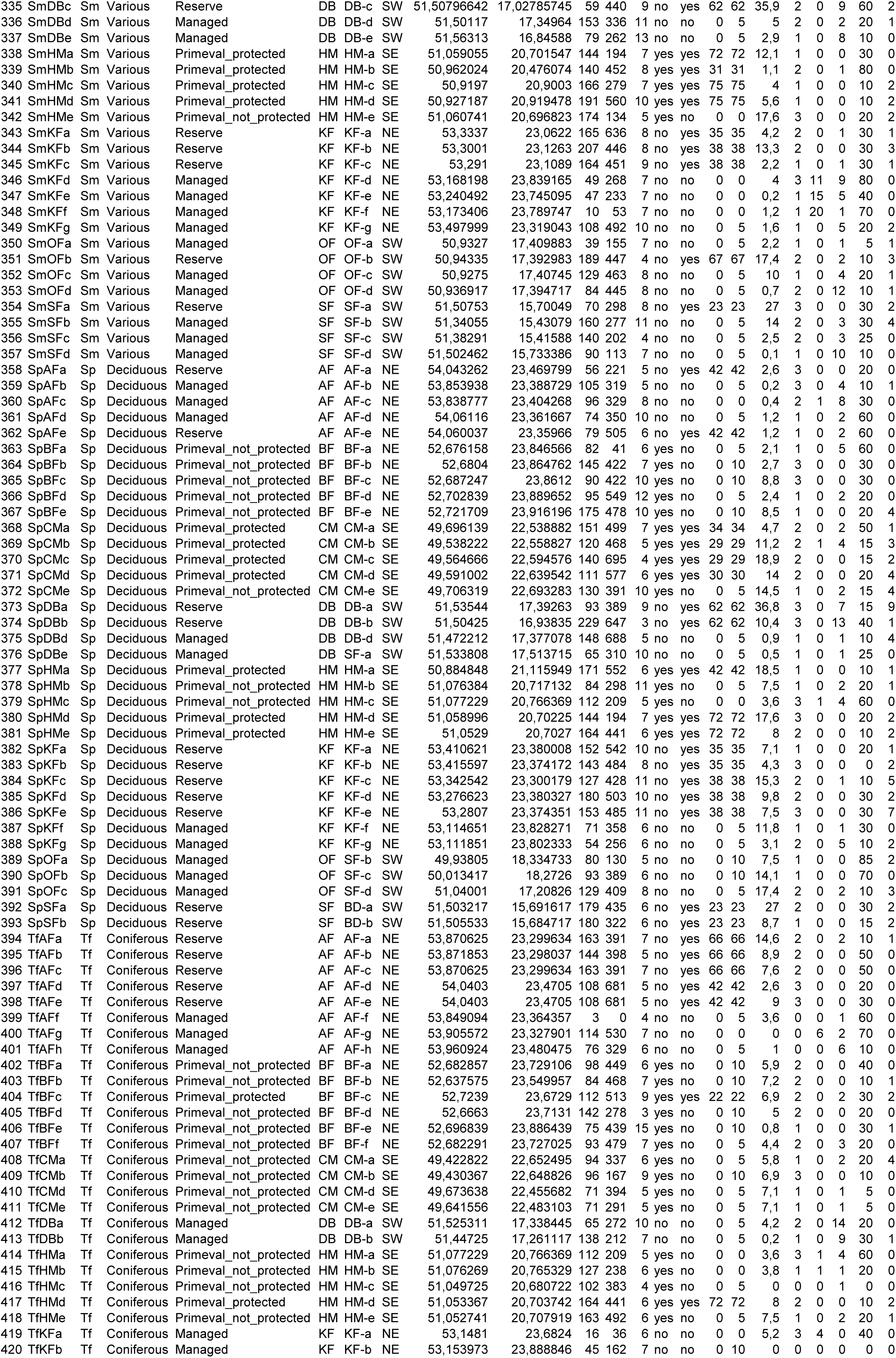
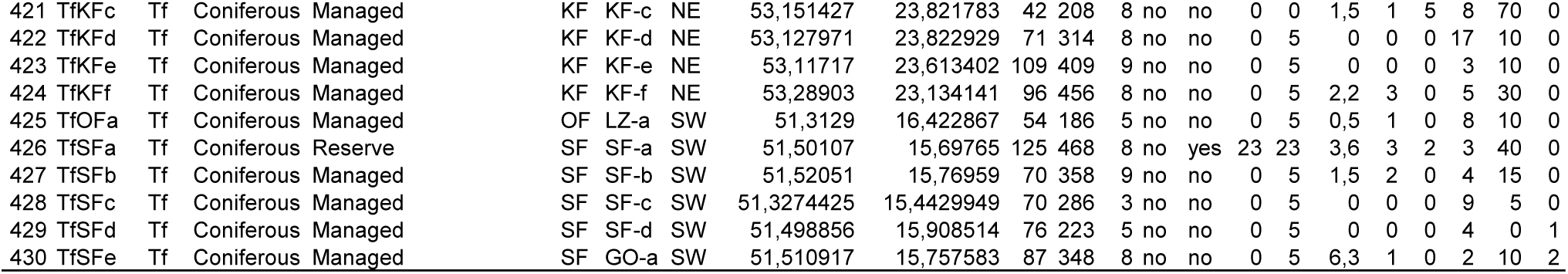
Characteristics of forest plots where saproxylic beetles were sampled for the study.

**Table S2.**
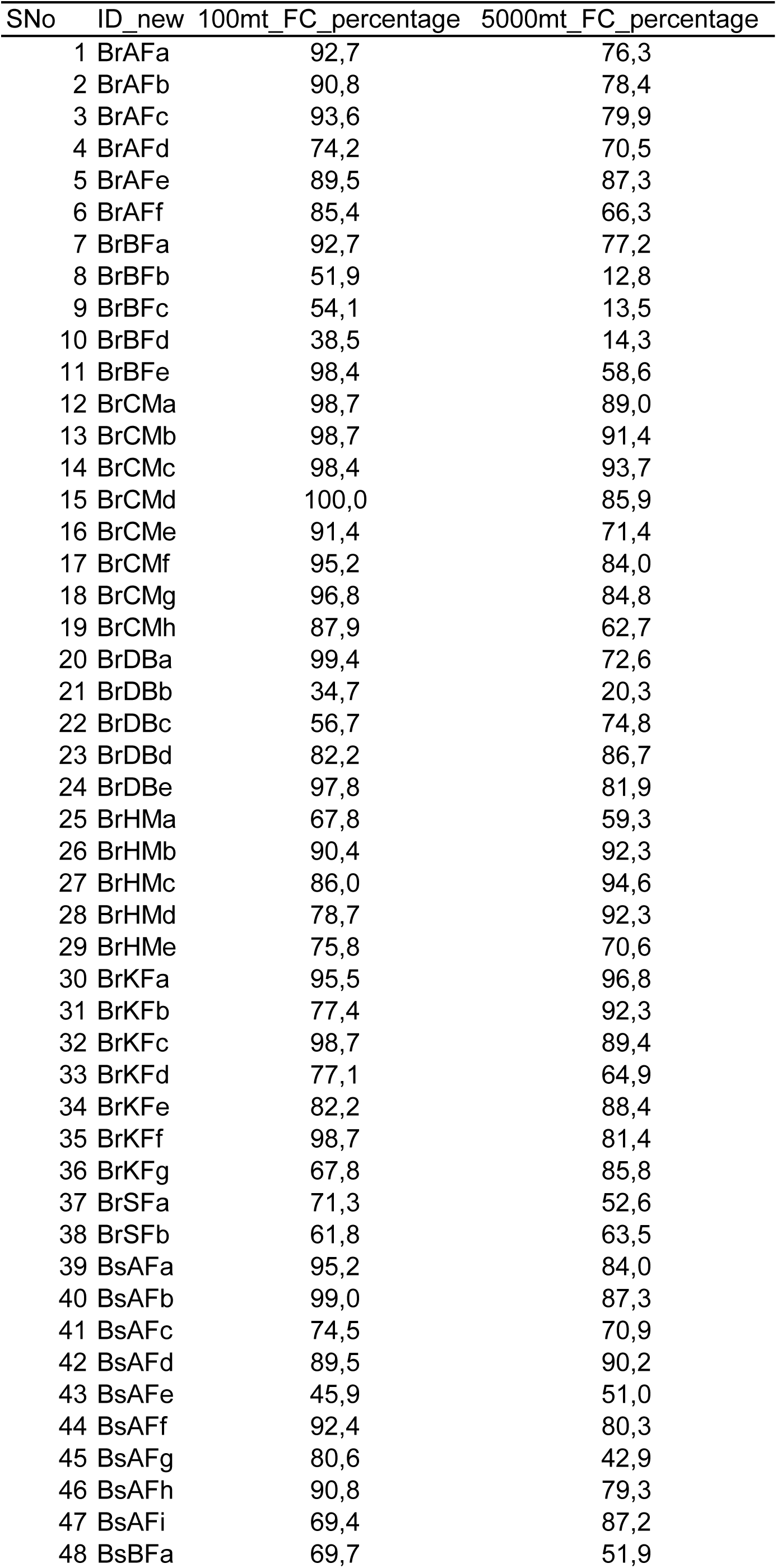
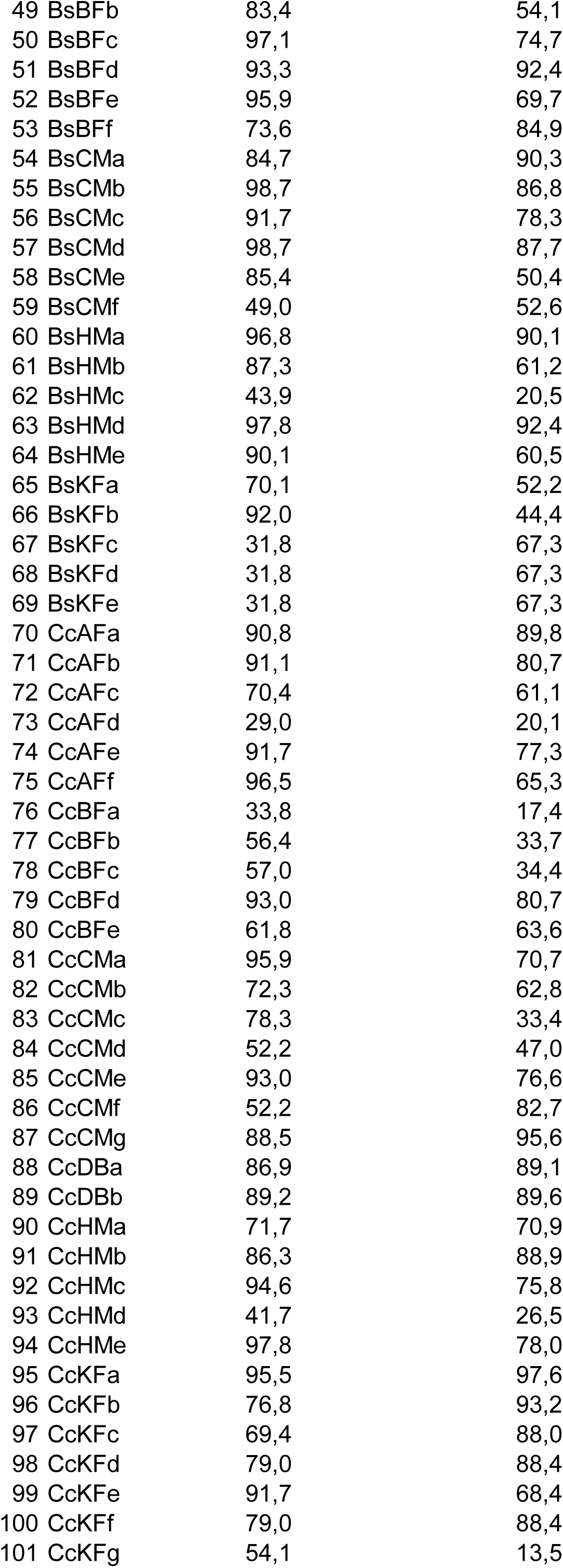
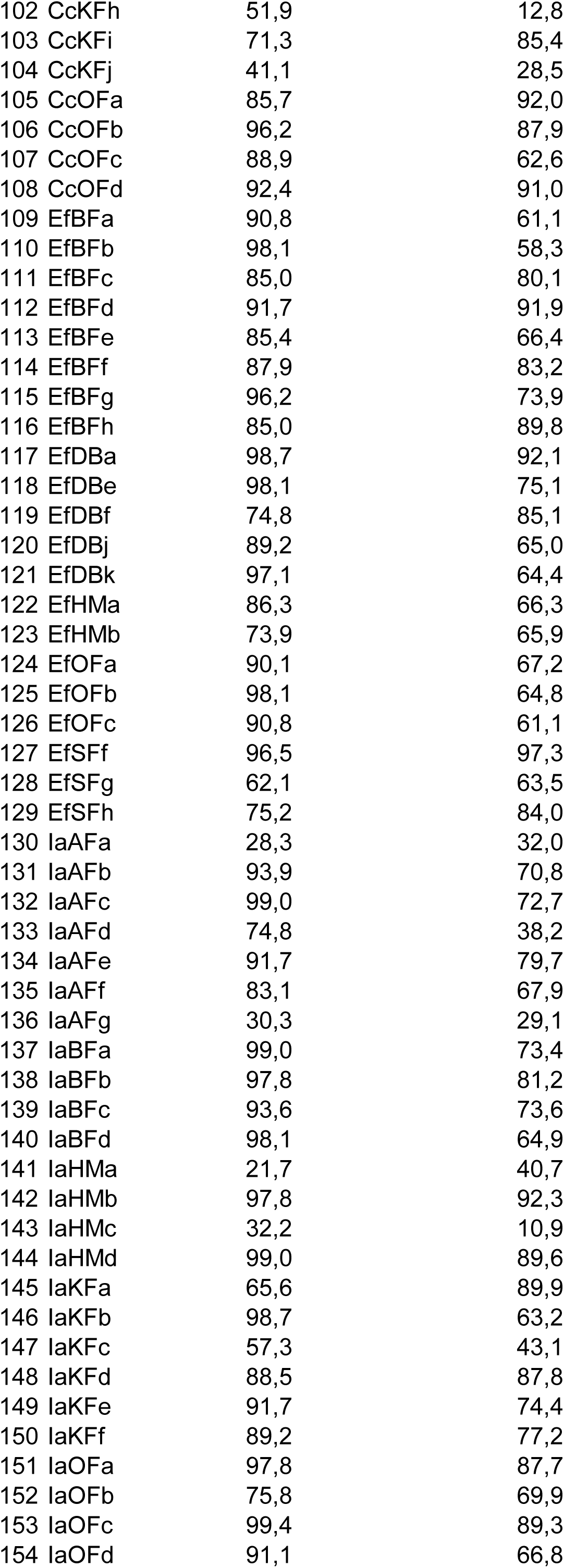
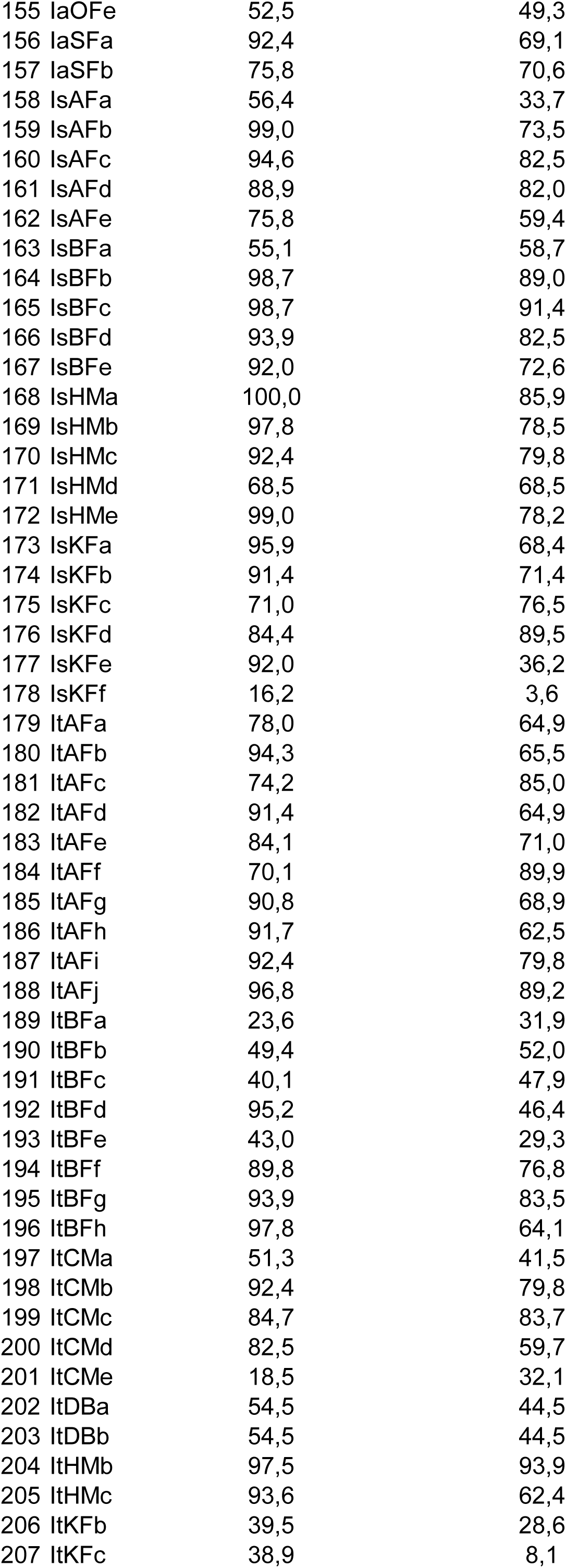
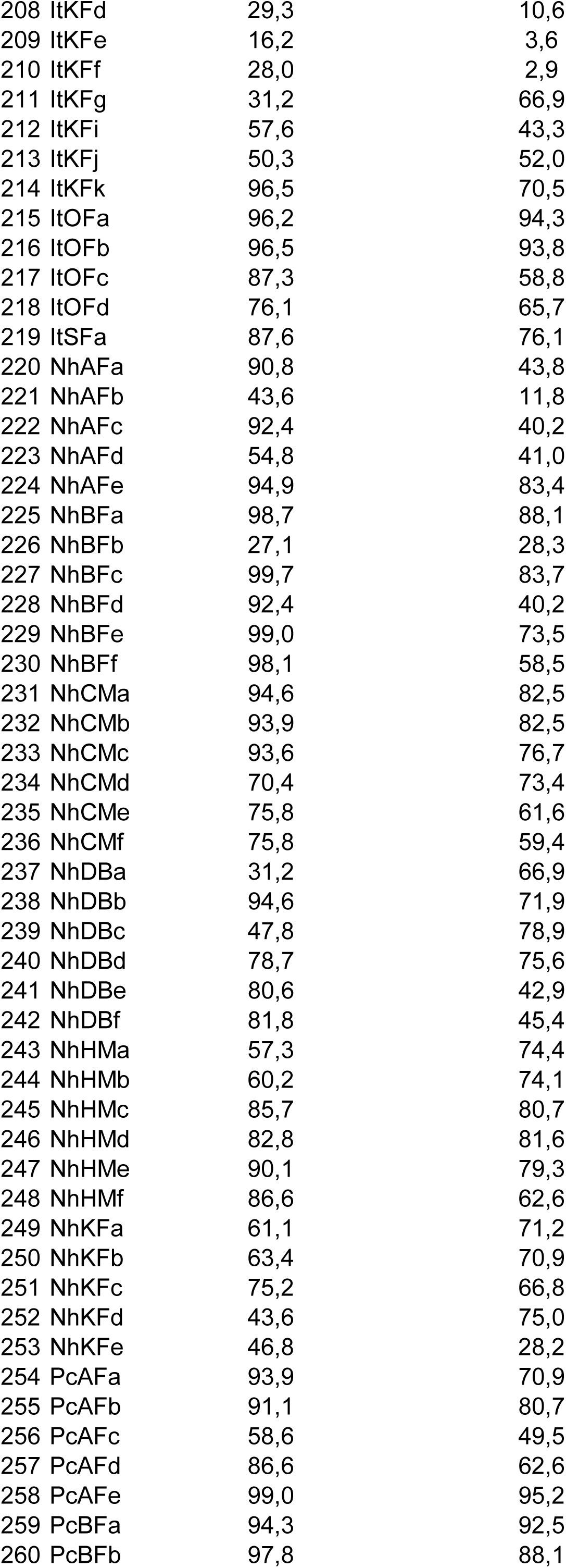
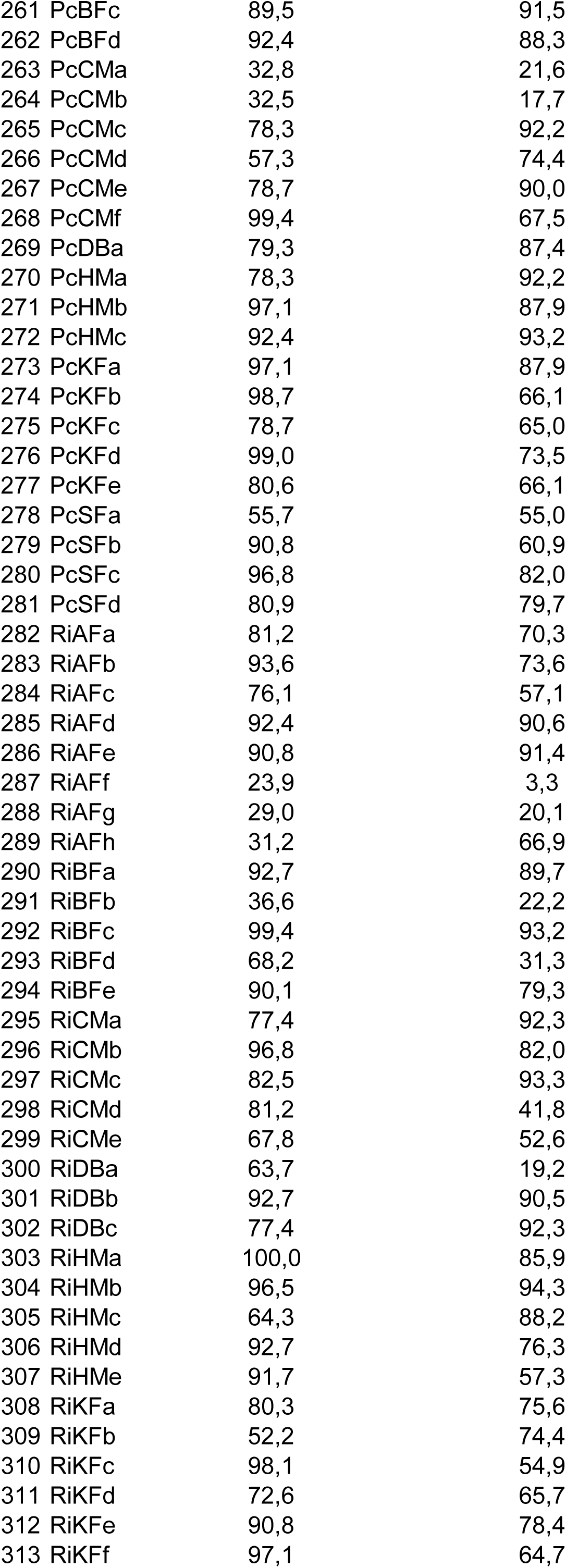
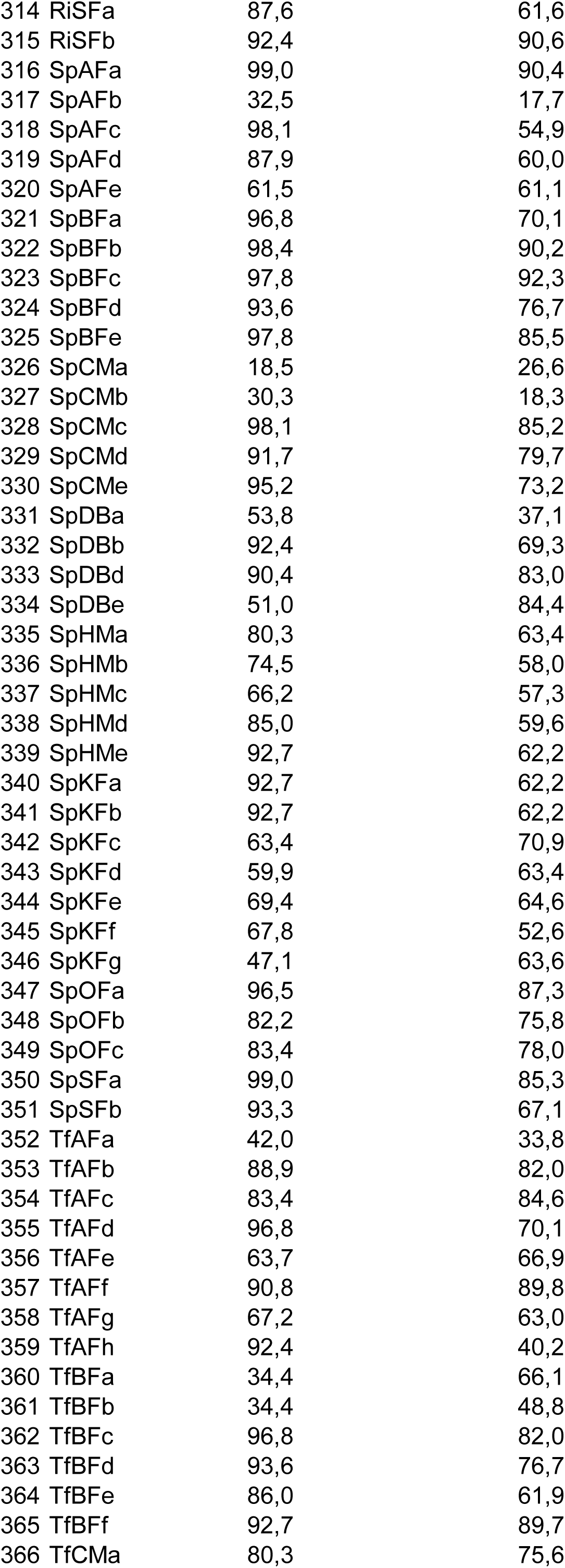
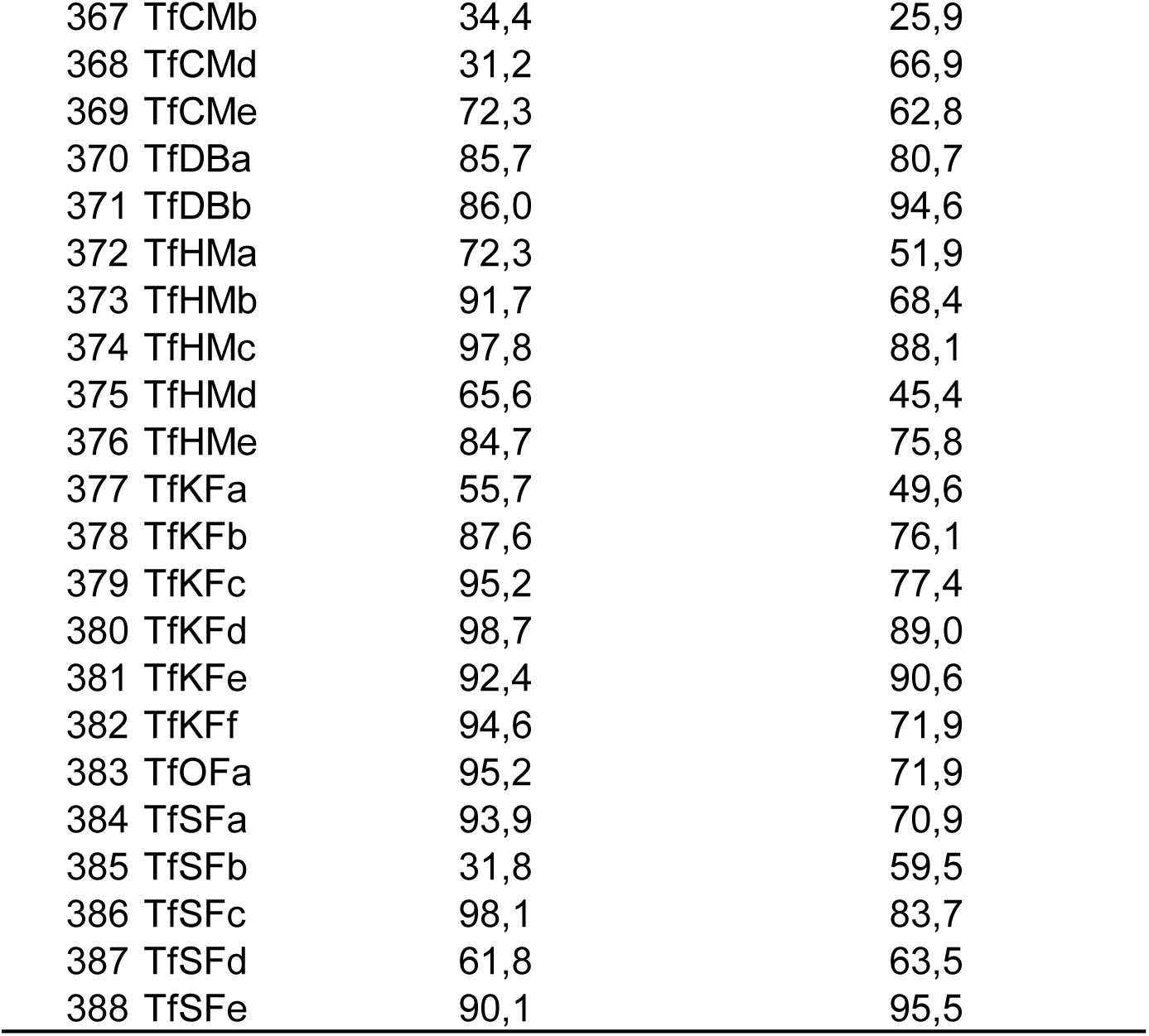
Summary of forest cover in 100m and 5000m buffers around examined plots where saproxylic beetles were sampled for the study.

| SNo | ID_new | 100mt_FC_percentage | 5000mt_FC_percentage |
| --- | --- | --- | --- |
| 1 | BrAFa | 92,7 | 76,3 |
| 2 | BrAFb | 90,8 | 78,4 |
| 3 | BrAFc | 93,6 | 79,9 |
| 4 | BrAFd | 74,2 | 70,5 |
| 5 | BrAFe | 89,5 | 87,3 |
| 6 | BrAFf | 85,4 | 66,3 |
| 7 | BrBFa | 92,7 | 77,2 |
| 8 | BrBFb | 51,9 | 12,8 |
| 9 | BrBFc | 54,1 | 13,5 |
| 10 | BrBFd | 38,5 | 14,3 |
| 11 | BrBFe | 98,4 | 58,6 |
| 12 | BrCMa | 98,7 | 89,0 |
| 13 | BrCMb | 98,7 | 91,4 |
| 14 | BrCMc | 98,4 | 93,7 |
| 15 | BrCMD | 100,0 | 85,9 |
| 16 | BrCMe | 91,4 | 71,4 |
| 17 | BrCMf | 95,2 | 84,0 |
| 18 | BrCMg | 96,8 | 84,8 |
| 19 | BrCMh | 87,9 | 62,7 |
| 20 | BrDBa | 99,4 | 72,6 |
| 21 | BrDBb | 34,7 | 20,3 |
| 22 | BrDBc | 56,7 | 74,8 |
| 23 | BrDBd | 82,2 | 86,7 |
| 24 | BrDBe | 97,8 | 81,9 |
| 25 | BrHMa | 67,8 | 59,3 |
| 26 | BrHMb | 90,4 | 92,3 |
| 27 | BrHMc | 86,0 | 94,6 |
| 28 | BrHMd | 78,7 | 92,3 |
| 29 | BrHMe | 75,8 | 70,6 |
| 30 | BrKF a | 95,5 | 96,8 |
| 31 | BrKF b | 77,4 | 92,3 |
| 32 | BrKF c | 98,7 | 89,4 |
| 33 | BrKF d | 77,1 | 64,9 |
| 34 | BrKF e | 82,2 | 88,4 |
| 35 | BrKF f | 98,7 | 81,4 |
| 36 | BrKF g | 67,8 | 85,8 |
| 37 | BrSF a | 71,3 | 52,6 |
| 38 | BrSF b | 61,8 | 63,5 |
| 39 | BsAF a | 95,2 | 84,0 |
| 40 | BsAF b | 99,0 | 87,3 |
| 41 | BsAF c | 74,5 | 70,9 |
| 42 | BsAF d | 89,5 | 90,2 |
| 43 | BsAF e | 45,9 | 51,0 |
| 44 | BsAF f | 92,4 | 80,3 |
| 45 | BsAF g | 80,6 | 42,9 |
| 46 | BsAF h | 90,8 | 79,3 |
| 47 | BsAF i | 69,4 | 87,2 |
| 48 | BsBF a | 69,7 | 51,9 |

|  |  |  |
| --- | --- | --- |
| 49 BsBFb | 83,4 | 54,1 |
| 50 BsBFc | 97,1 | 74,7 |
| 51 BsBFd | 93,3 | 92,4 |
| 52 BsBFfe | 95,9 | 69,7 |
| 53 BsBFf | 73,6 | 84,9 |
| 54 BsCMa | 84,7 | 90,3 |
| 55 BsCMb | 98,7 | 86,8 |
| 56 BsCMc | 91,7 | 78,3 |
| 57 BsCMD | 98,7 | 87,7 |
| 58 BsCMe | 85,4 | 50,4 |
| 59 BsCMf | 49,0 | 52,6 |
| 60 BsHMa | 96,8 | 90,1 |
| 61 BsHMb | 87,3 | 61,2 |
| 62 BsHMc | 43,9 | 20,5 |
| 63 BsHMD | 97,8 | 92,4 |
| 64 BsHMe | 90,1 | 60,5 |
| 65 BsKFa | 70,1 | 52,2 |
| 66 BsKFb | 92,0 | 44,4 |
| 67 BsKFc | 31,8 | 67,3 |
| 68 BsKFd | 31,8 | 67,3 |
| 69 BsKFe | 31,8 | 67,3 |
| 70 CcAFa | 90,8 | 89,8 |
| 71 CcAFb | 91,1 | 80,7 |
| 72 CcAFc | 70,4 | 61,1 |
| 73 CcAFd | 29,0 | 20,1 |
| 74 CcAFfe | 91,7 | 77,3 |
| 75 CcAFf | 96,5 | 65,3 |
| 76 CcBFa | 33,8 | 17,4 |
| 77 CcBFb | 56,4 | 33,7 |
| 78 CcBFc | 57,0 | 34,4 |
| 79 CcBFd | 93,0 | 80,7 |
| 80 CcBFfe | 61,8 | 63,6 |
| 81 CcCMA | 95,9 | 70,7 |
| 82 CcCMB | 72,3 | 62,8 |
| 83 CcCMc | 78,3 | 33,4 |
| 84 CcCMD | 52,2 | 47,0 |
| 85 CcCMe | 93,0 | 76,6 |
| 86 CcCMf | 52,2 | 82,7 |
| 87 CcCMg | 88,5 | 95,6 |
| 88 CcDBa | 86,9 | 89,1 |
| 89 CcDBb | 89,2 | 89,6 |
| 90 CcHMa | 71,7 | 70,9 |
| 91 CcHMb | 86,3 | 88,9 |
| 92 CcHMc | 94,6 | 75,8 |
| 93 CcHMD | 41,7 | 26,5 |
| 94 CcHMe | 97,8 | 78,0 |
| 95 CcKFa | 95,5 | 97,6 |
| 96 CcKFb | 76,8 | 93,2 |
| 97 CcKFc | 69,4 | 88,0 |
| 98 CcKFd | 79,0 | 88,4 |
| 99 CcKFe | 91,7 | 68,4 |
| 100 CcKFf | 79,0 | 88,4 |
| 101 CcKFg | 54,1 | 13,5 |
| 102 CcKFh | 51,9 | 12,8 |
| 103 CcKFi | 71,3 | 85,4 |
| 104 CcKFj | 41,1 | 28,5 |
| 105 CcOFa | 85,7 | 92,0 |
| 106 CcOFb | 96,2 | 87,9 |
| 107 CcOFc | 88,9 | 62,6 |
| 108 CcOFd | 92,4 | 91,0 |
| 109 EfBFa | 90,8 | 61,1 |
| 110 EfBFb | 98,1 | 58,3 |
| 111 EfBFc | 85,0 | 80,1 |
| 112 EfBFd | 91,7 | 91,9 |
| 113 EfBFe | 85,4 | 66,4 |
| 114 EfBFf | 87,9 | 83,2 |
| 115 EfBFg | 96,2 | 73,9 |
| 116 EfBFh | 85,0 | 89,8 |
| 117 EfDBa | 98,7 | 92,1 |
| 118 EfDBe | 98,1 | 75,1 |
| 119 EfDBf | 74,8 | 85,1 |
| 120 EfDBj | 89,2 | 65,0 |
| 121 EfDBk | 97,1 | 64,4 |
| 122 EfHMa | 86,3 | 66,3 |
| 123 EfHMb | 73,9 | 65,9 |
| 124 EfOFa | 90,1 | 67,2 |
| 125 EfOFb | 98,1 | 64,8 |
| 126 EfOFc | 90,8 | 61,1 |
| 127 EfSFf | 96,5 | 97,3 |
| 128 EfSFg | 62,1 | 63,5 |
| 129 EfSFh | 75,2 | 84,0 |
| 130 IaAFa | 28,3 | 32,0 |
| 131 IaAFb | 93,9 | 70,8 |
| 132 IaAFc | 99,0 | 72,7 |
| 133 IaAFd | 74,8 | 38,2 |
| 134 IaAFe | 91,7 | 79,7 |
| 135 IaAFf | 83,1 | 67,9 |
| 136 IaAFg | 30,3 | 29,1 |
| 137 IaBFa | 99,0 | 73,4 |
| 138 IaBFb | 97,8 | 81,2 |
| 139 IaBFc | 93,6 | 73,6 |
| 140 IaBFd | 98,1 | 64,9 |
| 141 IaHMa | 21,7 | 40,7 |
| 142 IaHMb | 97,8 | 92,3 |
| 143 IaHMc | 32,2 | 10,9 |
| 144 IaHMd | 99,0 | 89,6 |
| 145 IaKF a | 65,6 | 89,9 |
| 146 IaKFb | 98,7 | 63,2 |
| 147 IaKFc | 57,3 | 43,1 |
| 148 IaKFd | 88,5 | 87,8 |
| 149 IaKFe | 91,7 | 74,4 |
| 150 IaKFf | 89,2 | 77,2 |
| 151 IaOFa | 97,8 | 87,7 |
| 152 IaOFb | 75,8 | 69,9 |
| 153 IaOFc | 99,4 | 89,3 |
| 154 IaOFd | 91,1 | 66,8 |
| 155 | laOFe | 52,5 | 49,3 |
| 156 | laSFa | 92,4 | 69,1 |
| 157 | laSFb | 75,8 | 70,6 |
| 158 | IsAFa | 56,4 | 33,7 |
| 159 | IsAFb | 99,0 | 73,5 |
| 160 | IsAFc | 94,6 | 82,5 |
| 161 | IsAFd | 88,9 | 82,0 |
| 162 | IsAFe | 75,8 | 59,4 |
| 163 | IsBFa | 55,1 | 58,7 |
| 164 | IsBFb | 98,7 | 89,0 |
| 165 | IsBFc | 98,7 | 91,4 |
| 166 | IsBFd | 93,9 | 82,5 |
| 167 | IsBFe | 92,0 | 72,6 |
| 168 | IsHMa | 100,0 | 85,9 |
| 169 | IsHMb | 97,8 | 78,5 |
| 170 | IsHMc | 92,4 | 79,8 |
| 171 | IsHMd | 68,5 | 68,5 |
| 172 | IsHMe | 99,0 | 78,2 |
| 173 | IsKF a | 95,9 | 68,4 |
| 174 | IsKF b | 91,4 | 71,4 |
| 175 | IsKF c | 71,0 | 76,5 |
| 176 | IsKF d | 84,4 | 89,5 |
| 177 | IsKF e | 92,0 | 36,2 |
| 178 | IsKF f | 16,2 | 3,6 |
| 179 | ItAF a | 78,0 | 64,9 |
| 180 | ItAF b | 94,3 | 65,5 |
| 181 | ItAF c | 74,2 | 85,0 |
| 182 | ItAF d | 91,4 | 64,9 |
| 183 | ItAF e | 84,1 | 71,0 |
| 184 | ItAF f | 70,1 | 89,9 |
| 185 | ItAF g | 90,8 | 68,9 |
| 186 | ItAF h | 91,7 | 62,5 |
| 187 | ItAF i | 92,4 | 79,8 |
| 188 | ItAF j | 96,8 | 89,2 |
| 189 | ItBF a | 23,6 | 31,9 |
| 190 | ItBF b | 49,4 | 52,0 |
| 191 | ItBF c | 40,1 | 47,9 |
| 192 | ItBF d | 95,2 | 46,4 |
| 193 | ItBF e | 43,0 | 29,3 |
| 194 | ItBF f | 89,8 | 76,8 |
| 195 | ItBF g | 93,9 | 83,5 |
| 196 | ItBF h | 97,8 | 64,1 |
| 197 | ItCM a | 51,3 | 41,5 |
| 198 | ItCM b | 92,4 | 79,8 |
| 199 | ItCM c | 84,7 | 83,7 |
| 200 | ItCM d | 82,5 | 59,7 |
| 201 | ItCM e | 18,5 | 32,1 |
| 202 | ItDB a | 54,5 | 44,5 |
| 203 | ItDB b | 54,5 | 44,5 |
| 204 | ItHM b | 97,5 | 93,9 |
| 205 | ItHM c | 93,6 | 62,4 |
| 206 | ItKF b | 39,5 | 28,6 |
| 207 | ItKF c | 38,9 | 8,1 |
| 208 | ItKFd | 29,3 | 10,6 |
| 209 | ItKFe | 16,2 | 3,6 |
| 210 | ItKFf | 28,0 | 2,9 |
| 211 | ItKFg | 31,2 | 66,9 |
| 212 | ItKFi | 57,6 | 43,3 |
| 213 | ItKFj | 50,3 | 52,0 |
| 214 | ItKFk | 96,5 | 70,5 |
| 215 | ItOFa | 96,2 | 94,3 |
| 216 | ItOFb | 96,5 | 93,8 |
| 217 | ItOFc | 87,3 | 58,8 |
| 218 | ItOFd | 76,1 | 65,7 |
| 219 | ItSFa | 87,6 | 76,1 |
| 220 | NhAFa | 90,8 | 43,8 |
| 221 | NhAFb | 43,6 | 11,8 |
| 222 | NhAFc | 92,4 | 40,2 |
| 223 | NhAFd | 54,8 | 41,0 |
| 224 | NhAFe | 94,9 | 83,4 |
| 225 | NhBFa | 98,7 | 88,1 |
| 226 | NhBFb | 27,1 | 28,3 |
| 227 | NhBFc | 99,7 | 83,7 |
| 228 | NhBFd | 92,4 | 40,2 |
| 229 | NhBFe | 99,0 | 73,5 |
| 230 | NhBFf | 98,1 | 58,5 |
| 231 | NhCMa | 94,6 | 82,5 |
| 232 | NhCMb | 93,9 | 82,5 |
| 233 | NhCMc | 93,6 | 76,7 |
| 234 | NhCMD | 70,4 | 73,4 |
| 235 | NhCMe | 75,8 | 61,6 |
| 236 | NhCMf | 75,8 | 59,4 |
| 237 | NhDBa | 31,2 | 66,9 |
| 238 | NhDBb | 94,6 | 71,9 |
| 239 | NhDBc | 47,8 | 78,9 |
| 240 | NhDBd | 78,7 | 75,6 |
| 241 | NhDBe | 80,6 | 42,9 |
| 242 | NhDBf | 81,8 | 45,4 |
| 243 | NhHMa | 57,3 | 74,4 |
| 244 | NhHMb | 60,2 | 74,1 |
| 245 | NhHMc | 85,7 | 80,7 |
| 246 | NhHMd | 82,8 | 81,6 |
| 247 | NhHMe | 90,1 | 79,3 |
| 248 | NhHMf | 86,6 | 62,6 |
| 249 | NhKF a | 61,1 | 71,2 |
| 250 | NhKF b | 63,4 | 70,9 |
| 251 | NhKF c | 75,2 | 66,8 |
| 252 | NhKF d | 43,6 | 75,0 |
| 253 | NhKF e | 46,8 | 28,2 |
| 254 | PcAFa | 93,9 | 70,9 |
| 255 | PcAFb | 91,1 | 80,7 |
| 256 | PcAFc | 58,6 | 49,5 |
| 257 | PcAFd | 86,6 | 62,6 |
| 258 | PcAF e | 99,0 | 95,2 |
| 259 | PcBFa | 94,3 | 92,5 |
| 260 | PcBFb | 97,8 | 88,1 |

|  |  |  |
| --- | --- | --- |
| 261 PcBFc | 89,5 | 91,5 |
| 262 PcBFd | 92,4 | 88,3 |
| 263 PcCMa | 32,8 | 21,6 |
| 264 PcCMb | 32,5 | 17,7 |
| 265 PcCMc | 78,3 | 92,2 |
| 266 PcCMD | 57,3 | 74,4 |
| 267 PcCMe | 78,7 | 90,0 |
| 268 PcCMf | 99,4 | 67,5 |
| 269 PcDBa | 79,3 | 87,4 |
| 270 PcHMa | 78,3 | 92,2 |
| 271 PcHMb | 97,1 | 87,9 |
| 272 PcHMc | 92,4 | 93,2 |
| 273 PcKF<sub>a</sub> | 97,1 | 87,9 |
| 274 PcKF<sub>b</sub> | 98,7 | 66,1 |
| 275 PcKF<sub>c</sub> | 78,7 | 65,0 |
| 276 PcKF<sub>d</sub> | 99,0 | 73,5 |
| 277 PcKF<sub>e</sub> | 80,6 | 66,1 |
| 278 PcSF<sub>a</sub> | 55,7 | 55,0 |
| 279 PcSF<sub>b</sub> | 90,8 | 60,9 |
| 280 PcSF<sub>c</sub> | 96,8 | 82,0 |
| 281 PcSF<sub>d</sub> | 80,9 | 79,7 |
| 282 RiAF<sub>a</sub> | 81,2 | 70,3 |
| 283 RiAF<sub>b</sub> | 93,6 | 73,6 |
| 284 RiAF<sub>c</sub> | 76,1 | 57,1 |
| 285 RiAF<sub>d</sub> | 92,4 | 90,6 |
| 286 RiAF<sub>e</sub> | 90,8 | 91,4 |
| 287 RiAF<sub>f</sub> | 23,9 | 3,3 |
| 288 RiAF<sub>g</sub> | 29,0 | 20,1 |
| 289 RiAF<sub>h</sub> | 31,2 | 66,9 |
| 290 RiBF<sub>a</sub> | 92,7 | 89,7 |
| 291 RiBF<sub>b</sub> | 36,6 | 22,2 |
| 292 RiBF<sub>c</sub> | 99,4 | 93,2 |
| 293 RiBF<sub>d</sub> | 68,2 | 31,3 |
| 294 RiBF<sub>e</sub> | 90,1 | 79,3 |
| 295 RiCM<sub>a</sub> | 77,4 | 92,3 |
| 296 RiCM<sub>b</sub> | 96,8 | 82,0 |
| 297 RiCM<sub>c</sub> | 82,5 | 93,3 |
| 298 RiCM<sub>d</sub> | 81,2 | 41,8 |
| 299 RiCM<sub>e</sub> | 67,8 | 52,6 |
| 300 RiDB<sub>a</sub> | 63,7 | 19,2 |
| 301 RiDB<sub>b</sub> | 92,7 | 90,5 |
| 302 RiDB<sub>c</sub> | 77,4 | 92,3 |
| 303 RiHMa | 100,0 | 85,9 |
| 304 RiHMb | 96,5 | 94,3 |
| 305 RiHMc | 64,3 | 88,2 |
| 306 RiHMD | 92,7 | 76,3 |
| 307 RiHMe | 91,7 | 57,3 |
| 308 RiKF<sub>a</sub> | 80,3 | 75,6 |
| 309 RiKF<sub>b</sub> | 52,2 | 74,4 |
| 310 RiKF<sub>c</sub> | 98,1 | 54,9 |
| 311 RiKF<sub>d</sub> | 72,6 | 65,7 |
| 312 RiKF<sub>e</sub> | 90,8 | 78,4 |
| 313 RiKF<sub>f</sub> | 97,1 | 64,7 |
| 314 RiSFa | 87,6 | 61,6 |
| 315 RiSFb | 92,4 | 90,6 |
| 316 SpAFa | 99,0 | 90,4 |
| 317 SpAFb | 32,5 | 17,7 |
| 318 SpAFc | 98,1 | 54,9 |
| 319 SpAFd | 87,9 | 60,0 |
| 320 SpAFe | 61,5 | 61,1 |
| 321 SpBFa | 96,8 | 70,1 |
| 322 SpBFb | 98,4 | 90,2 |
| 323 SpBFc | 97,8 | 92,3 |
| 324 SpBFd | 93,6 | 76,7 |
| 325 SpBFfe | 97,8 | 85,5 |
| 326 SpCMa | 18,5 | 26,6 |
| 327 SpCMb | 30,3 | 18,3 |
| 328 SpCMc | 98,1 | 85,2 |
| 329 SpCMD | 91,7 | 79,7 |
| 330 SpCMe | 95,2 | 73,2 |
| 331 SpDBa | 53,8 | 37,1 |
| 332 SpDBb | 92,4 | 69,3 |
| 333 SpDBd | 90,4 | 83,0 |
| 334 SpDBe | 51,0 | 84,4 |
| 335 SpHMa | 80,3 | 63,4 |
| 336 SpHMb | 74,5 | 58,0 |
| 337 SpHMc | 66,2 | 57,3 |
| 338 SpHMD | 85,0 | 59,6 |
| 339 SpHMe | 92,7 | 62,2 |
| 340 SpKFa | 92,7 | 62,2 |
| 341 SpKFb | 92,7 | 62,2 |
| 342 SpKFc | 63,4 | 70,9 |
| 343 SpKFd | 59,9 | 63,4 |
| 344 SpKFe | 69,4 | 64,6 |
| 345 SpKff | 67,8 | 52,6 |
| 346 SpKFg | 47,1 | 63,6 |
| 347 SpOFa | 96,5 | 87,3 |
| 348 SpOFb | 82,2 | 75,8 |
| 349 SpOFc | 83,4 | 78,0 |
| 350 SpSFa | 99,0 | 85,3 |
| 351 SpSFb | 93,3 | 67,1 |
| 352 TfAFa | 42,0 | 33,8 |
| 353 TfAFb | 88,9 | 82,0 |
| 354 TfAFc | 83,4 | 84,6 |
| 355 TfAFd | 96,8 | 70,1 |
| 356 TfAFe | 63,7 | 66,9 |
| 357 TfAFf | 90,8 | 89,8 |
| 358 TfAFg | 67,2 | 63,0 |
| 359 TfAFh | 92,4 | 40,2 |
| 360 TfBFa | 34,4 | 66,1 |
| 361 TfBFb | 34,4 | 48,8 |
| 362 TfBFc | 96,8 | 82,0 |
| 363 TfBFd | 93,6 | 76,7 |
| 364 TfBFfe | 86,0 | 61,9 |
| 365 TfBFf | 92,7 | 89,7 |
| 366 TfCMa | 80,3 | 75,6 |
| 367 TfCMb | 34,4 | 25,9 |
| 368 TfCMd | 31,2 | 66,9 |
| 369 TfCMe | 72,3 | 62,8 |
| 370 TfDBa | 85,7 | 80,7 |
| 371 TfDBb | 86,0 | 94,6 |
| 372 TfHMa | 72,3 | 51,9 |
| 373 TfHMb | 91,7 | 68,4 |
| 374 TfHMc | 97,8 | 88,1 |
| 375 TfHMd | 65,6 | 45,4 |
| 376 TfHMe | 84,7 | 75,8 |
| 377 TfKF a | 55,7 | 49,6 |
| 378 TfKF b | 87,6 | 76,1 |
| 379 TfKF c | 95,2 | 77,4 |
| 380 TfKF d | 98,7 | 89,0 |
| 381 TfKF e | 92,4 | 90,6 |
| 382 TfKF f | 94,6 | 71,9 |
| 383 TfOF a | 95,2 | 71,9 |
| 384 TfSF a | 93,9 | 70,9 |
| 385 TfSF b | 31,8 | 59,5 |
| 386 TfSF c | 98,1 | 83,7 |
| 387 TfSF d | 61,8 | 63,5 |
| 388 TfSF e | 90,1 | 95,5 |

**Table S3.**
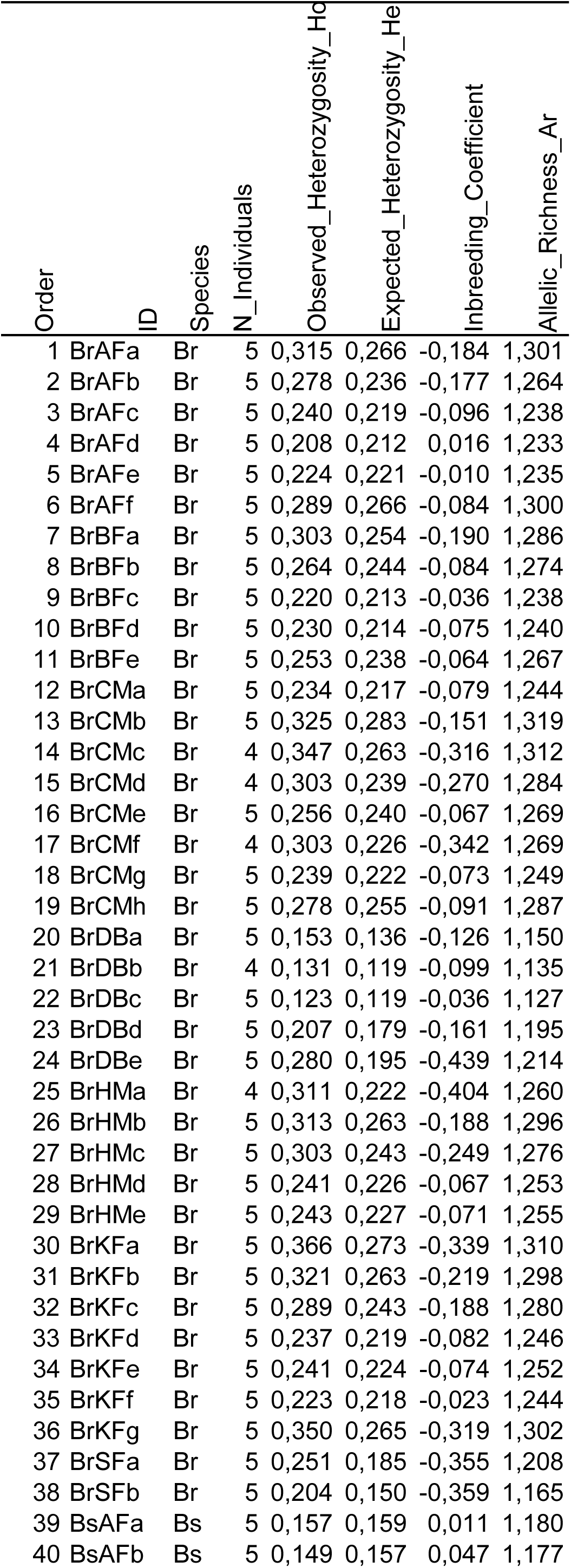
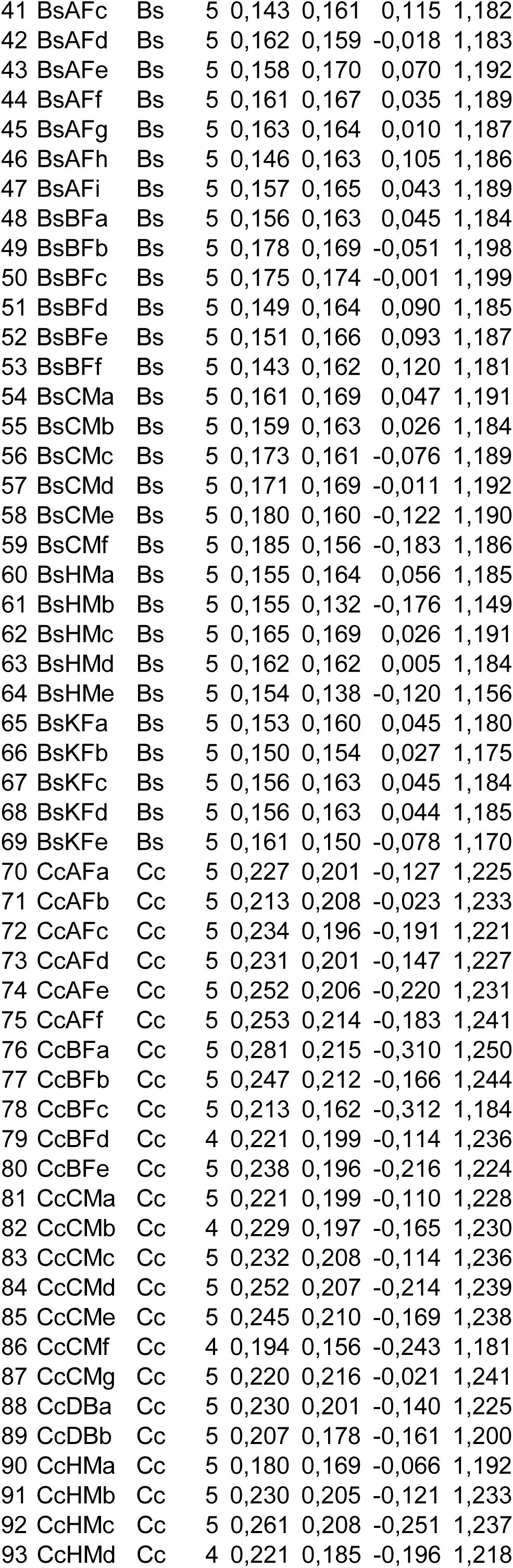
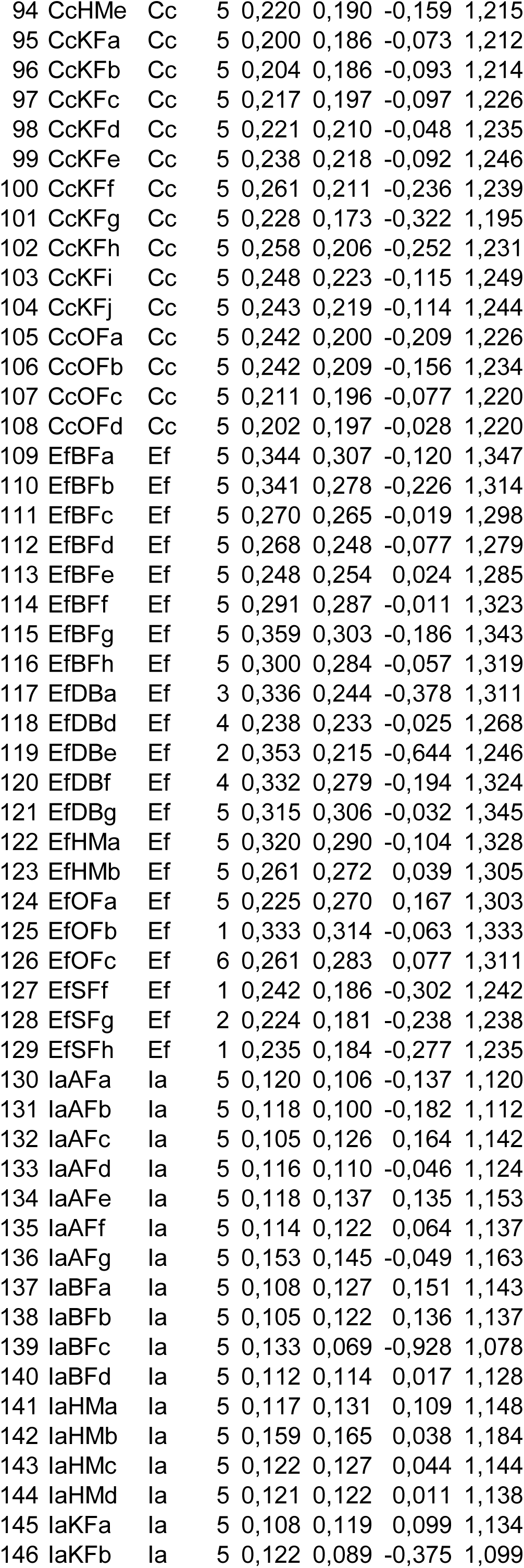
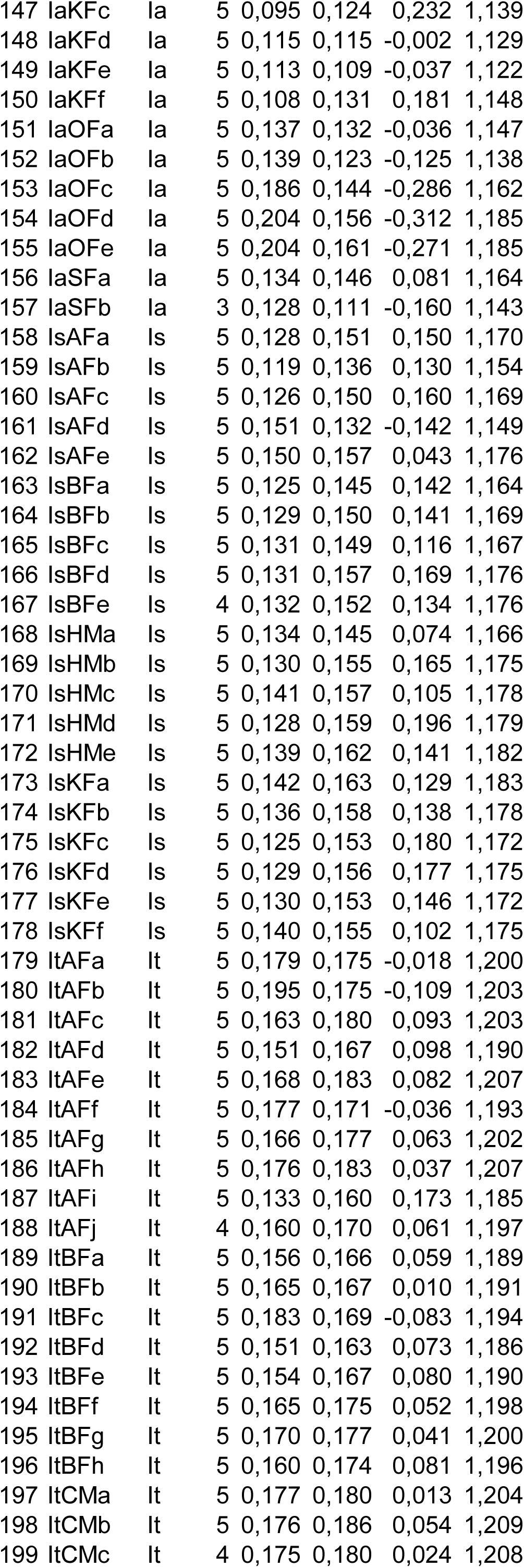
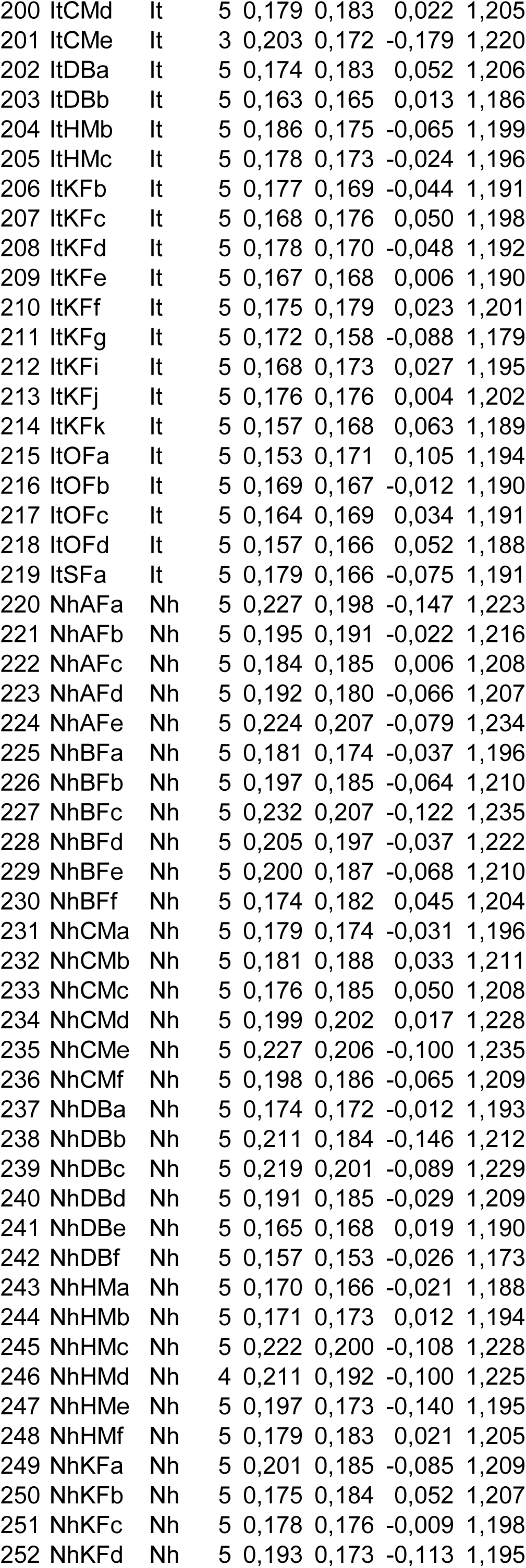
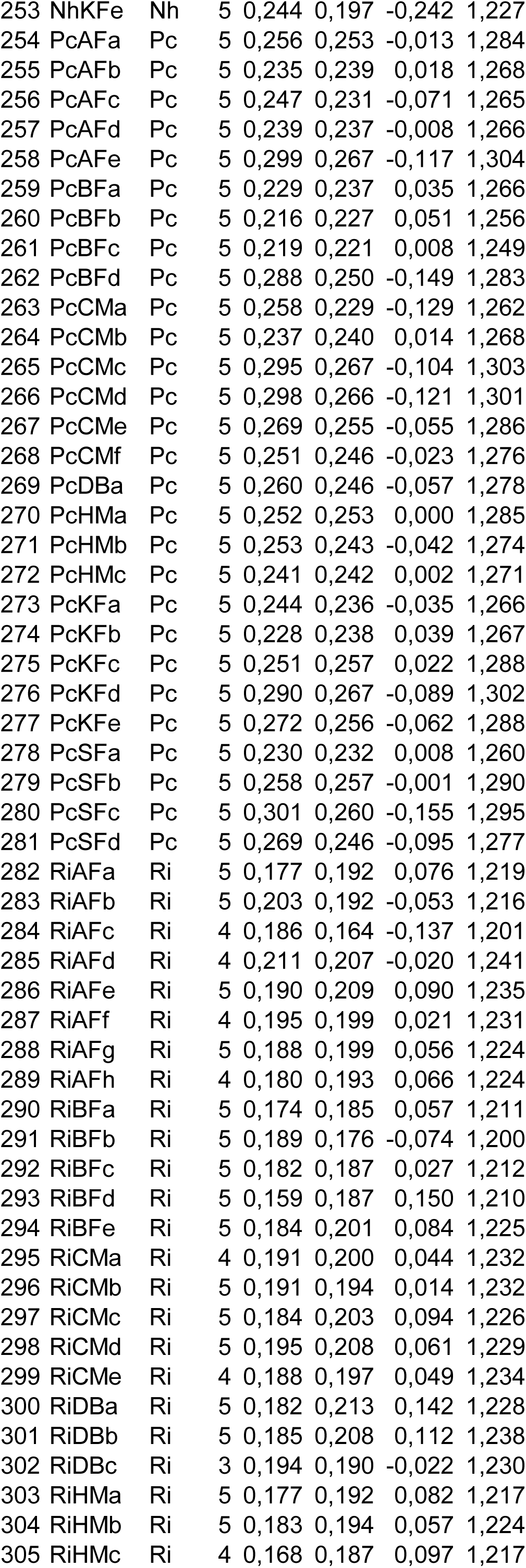
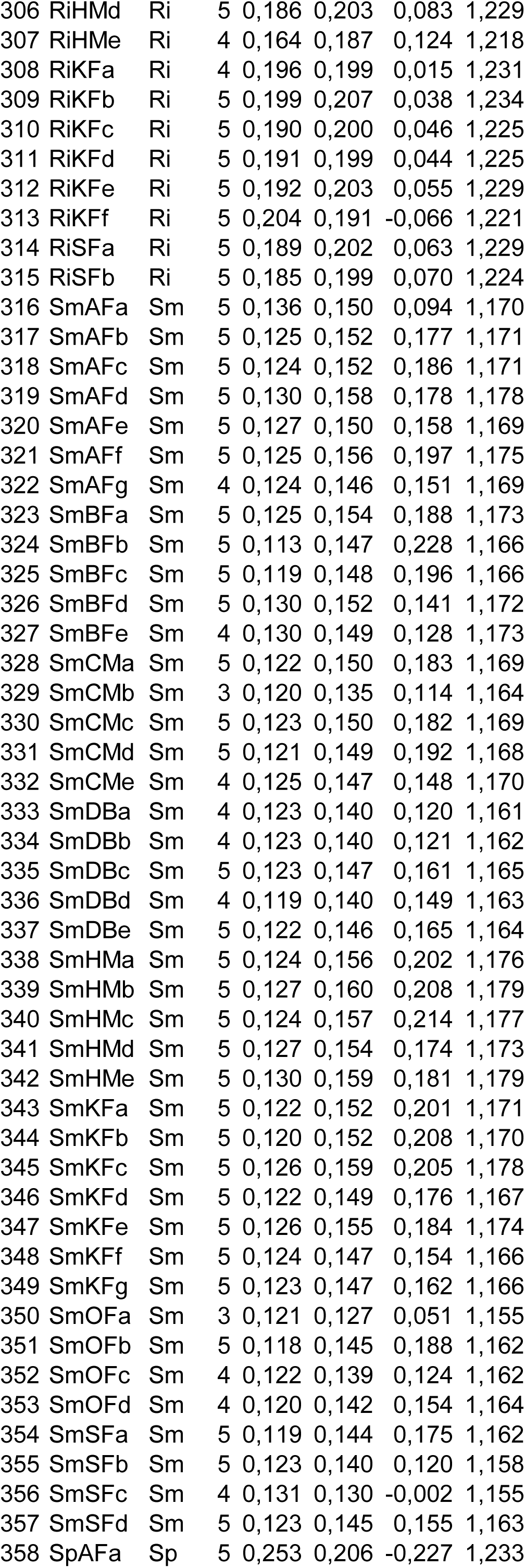
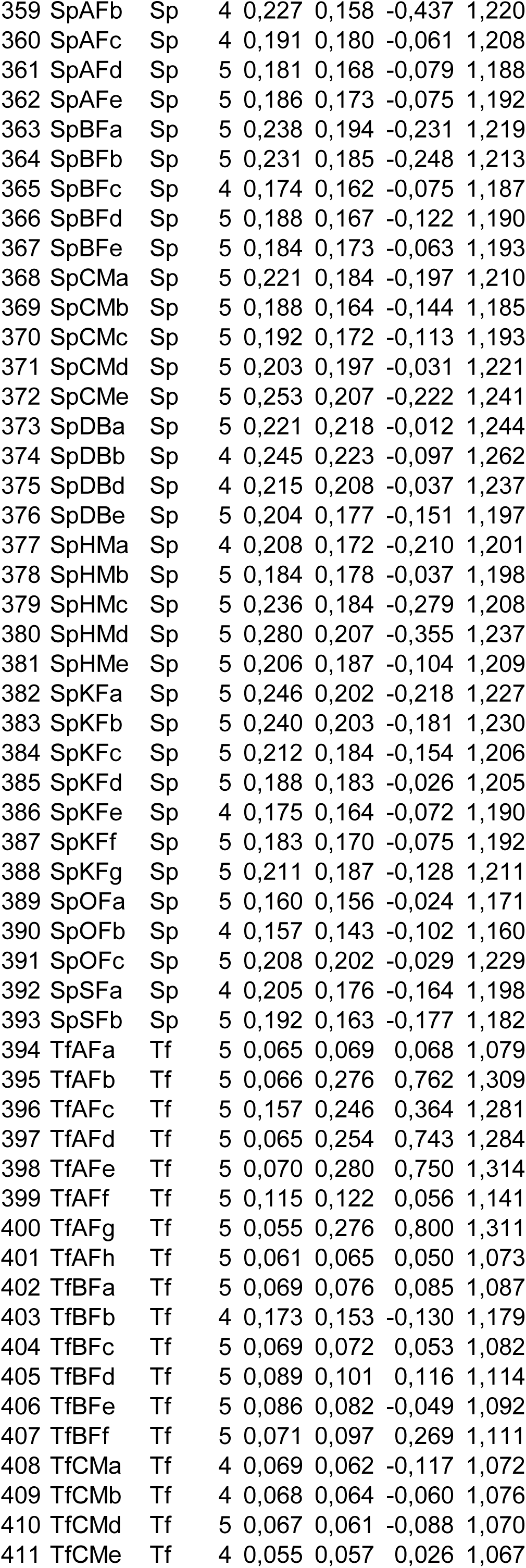
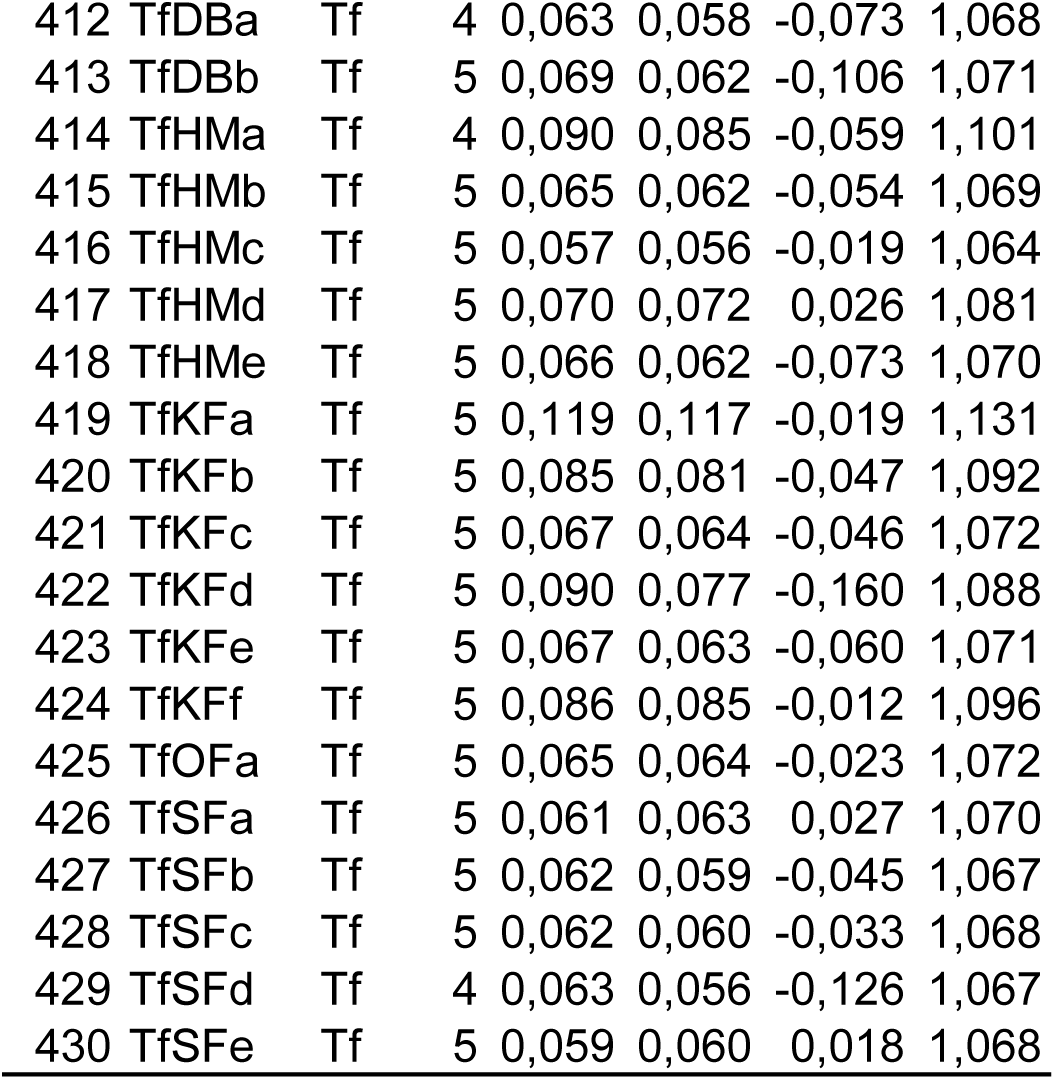
Characteristics of genetic metrics of saproxylic beetles sampled in selected plots.

